# Hippocampal place cells use vector computations to navigate

**DOI:** 10.1101/2021.06.23.449621

**Authors:** Jake Ormond, John O’Keefe

**Author notes:** corresponding authors Correspondence and requests for materials should be addressed to J. O., and J. O. K.

## Abstract

One function of the Hippocampal Cognitive Map is to provide information about salient locations in familiar environments such as those containing reward or danger, and to support navigation towards or away from those locations^1^. Although much is known about how the hippocampus encodes location in world-centred coordinates, how it supports flexible navigation is less well understood. We recorded from CA1 place cells while rats navigated to a goal or freely foraged on the honeycomb maze^2^. The maze tests the animal’s ability to navigate using indirect as well as direct paths to the goal and allows the directionality of place cells to be assessed at each choice point during traversal to the goal. Place fields showed strong directional polarization in the navigation task, and to a lesser extent during random foraging. This polarization was characterized by vector fields which converged to sinks distributed throughout the environment. The distribution of these *convergence sinks* was centred near the goal location, and the population vector field converged on the goal, providing a strong navigational signal. Changing the goal location led to the movement of *ConSinks* and vector fields towards the new goal and within-days, the ConSink distance to the goal decreased with continued training. The honeycomb maze allows the independent assessment of spatial representation and spatial action in place cell activity and shows how the latter depends on the former. The results suggest a vector-based model of how the hippocampus supports flexible navigation, allowing animals to select optimal paths to destinations from any location in the environment.

---

There is substantial evidence that the hippocampus acts as a cognitive map providing information about an animal’s current and future locations and how to navigate between them^1^. Lesions of the hippocampal formation reduce an animal’s ability to navigate to remembered locations, such as the escape platform in the Morris watermaze^3^. One strong candidate for underpinning navigation are the CA1 place cells which provide information about the animal’s current location^4^. In open-field foraging tasks lacking a specific goal, place cells provide an omnidirectional measure of current position^5,6^ (but see ^7^). When a goal is introduced, place cells become directional^8^ and their fields move in the direction of the goal when it is moved^9^. Recent work has shown that cells in the dorsal hippocampus encode heading towards a goal and other locations in the bat^10^, mouse7 and human^11^. However, an important caveat about such studies is that, once a goal is introduced, the animal usually moves towards the goal whenever possible, precluding assessment of the neuronal activity in non-goalward directions and locations.

In contrast, while navigating on the honeycomb maze^2^, the animal approaches a goal platform by a succession of binary choices between intermediate platforms, choosing the most efficient path towards the goal even when direct paths are not available (Fig. 1a, b; Extended Data 1a, b). Importantly, while the animal waits on each platform to make its next choice, it frequently scans around the platform perimeter, sampling the full range of possible headings (Fig. 1d left, Extended Data Fig. 2, Extended Data Video 1) permitting a veridical assessment of cell firing directionality.

**Figure 1.**
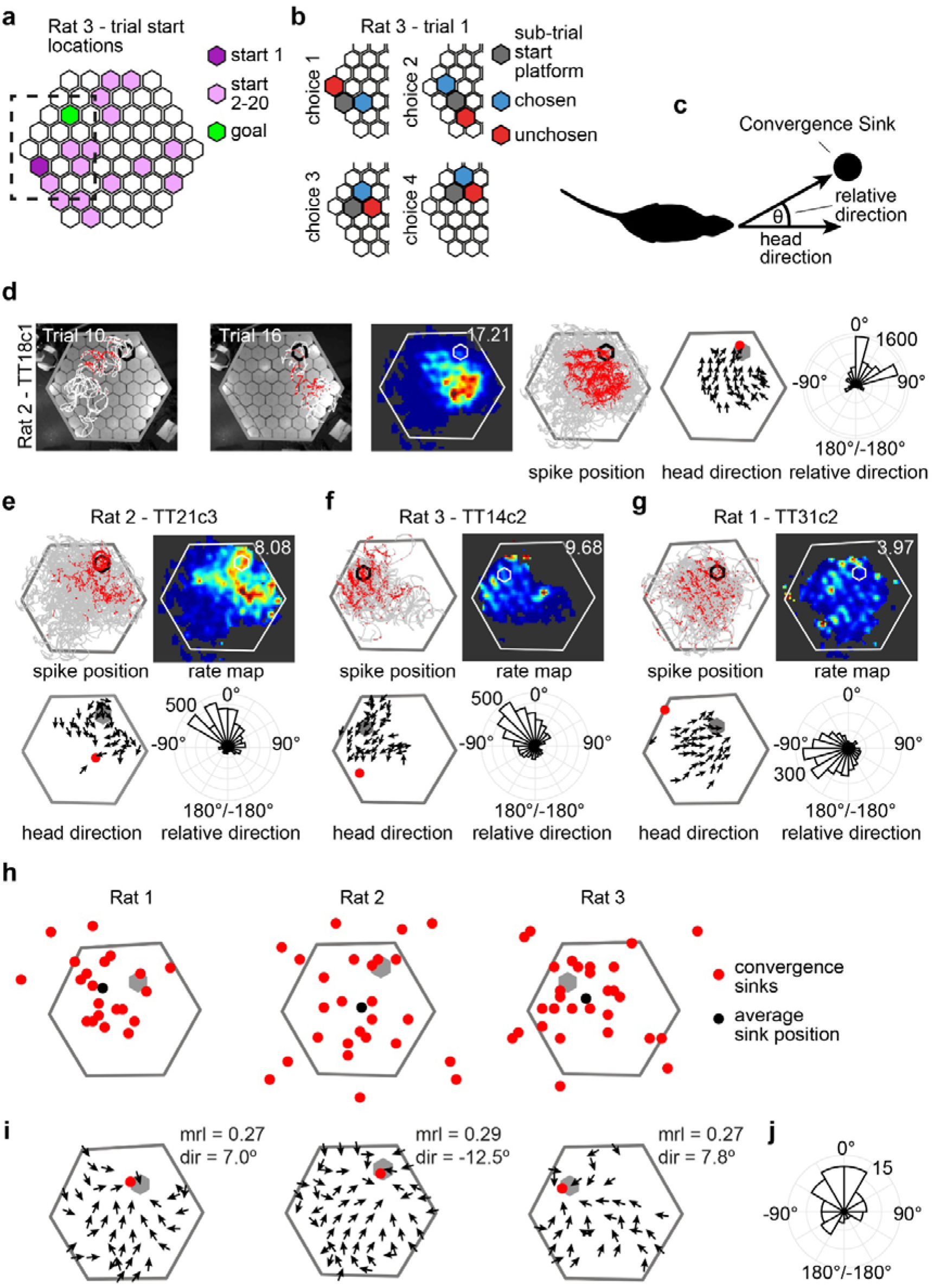
ConSinks and vector fields organize place cell activity during navigation on the honeycomb maze. **a**, Schematic of the honeycomb maze showing all start platforms and the goal platform from Rat 3’s session. Dashed box refers to the portion of the maze shown in (**b**). **b**, Schematic of the 4 choices that comprise trial 1 for Rat 3. Only the coloured platforms are raised. The animal starts each choice on the “sub-trial start” platform (grey) where it is confined until two adjacent platforms are raised (red and blue) and it makes its final choice by moving onto the “chosen” (blue) platform. **c**, Schematic showing how the animal’s heading direction relative to a reference point called the ConSink is calculated. The angle between the straight ahead head direction (0°) and the direction of the ConSink in egocentric space is called the relative direction. **d**, Representative example of a ConSink place cell. Left 2 panels, paths (white) and spikes (red) fired during 2 individual trials of the task. Perimeter of the goal platform in black. Middle 2 panels, place field heat map derived from spikes (in red) and dwell times (in grey) fired during the task plotted over the animal’s path. Note the scalloped firing patterns. Second from right, vector field depicts mean head direction at binned spatial positions. The ConSink for this cell is depicted as a filled red circle. Right, Polar plot showing the distribution of spike-associated head directions relative to the ConSink. **e**-**g**, Additional examples as in (**d**). **h**, ConSinks (in red) for all significant cells recorded during the task are widely distributed across the maze, and some are also located past the maze perimeter. ConSink centroids in black. **i**, Average vector fields for each of the three animals. “mrl”, mean resultant length; “dir”, mean relative direction. **j**, Mean relative directions of all significant ConSinks were non-uniformly distributed (*p* < 0.001, Rayleigh test), with a mean direction of −19° (not significantly different from 0°, one-sample test for mean angle).

## Place Cell Firing is organised by ConSinks

We recorded 266 CA1 place cells (defined as carrying significant spatial information^12^; Extended Data Fig. 3) from 3 rats (Rat 1: 89 cells; Rat 2: 94 cells; Rat 3: 83 cells) which had successfully learned the navigation task (Extended Data Fig. 1c-g). Of these, 77 (29%) displayed firing patterns within the place field which were best described by vector fields converging on a location that, following vector field notation, we term a convergence sink (ConSink; Fig. 1c-g). Each ConSink represents a location in allocentric co-ordinates relative to which the animal must maintain an egocentric relationship (Extended Data Fig 4). For example, the cell shown in Fig. 1d fired maximally when the animal’s head was pointing between 0° & 75° to the left of the ConSink located close to the goal (red circle). Often, the optimal bearing to the ConSink was located within a cone of 45° in front of the animal (Fig 1d-f), but for many it could be 90° or greater, to the side or even behind (e.g. Fig 1g). While ConSinks were scattered around the environment both on and off the maze, they were densest around the goal (Fig. 1h, Extended Data Fig. 5a). Importantly, the average of the vector fields for each animal converged to a population ConSink close to the goal (Fig 1i) and the average heading direction was in front of the animal, −19.2°, not significantly different from 0° (Fig 1j). ConSink tuning was stronger in every ConSink cell than head-direction tuning (Extended Data Fig. 5b).

## The population of ConSinks point to the goal

If ConSink directional firing supports navigation to a goal, their distribution might reflect this in 2 different ways. First, when the goal is shifted, they should move towards the new goal; second, with continued training to a fixed goal, the population of ConSinks might cluster closer around the goal.

We tested the goal shift prediction by running the same 3 animals in a goal shift experiment on a subsequent day (282 CA1 place cells; Rat 1: 93 cells; Rat 2: 96 cells; Rat 3: 93 cells): 13 trials with a familiar goal were followed, after some intermediary training (see Methods), by another 13 trials to a new goal. All 3 animals successfully learned the new goal location (Extended Data Fig. 6) and 32% (83/256) of principal cells during the original and 25% (70/275) during the shifted-goal navigations had significant ConSink tuning (Extended Data Fig. 7). Before the goal shift, ConSinks were organized around the original goal, as indicated by significantly shorter distances to it than to the new goal (Fig. 2a, b); after the switch, this was reversed (Fig. 2c, d), indicating that ConSinks were under the influence of goal location. While this reorganization primarily involved the substitution of new ConSinks for old, 25 cells with ConSinks during goal 1 continued to have ConSinks after the switch to goal 2 and the majority of these (16/25, 64%) moved in the direction of the new goal (Fig. 2e, f). Similarly, the vector fields and their associated population ConSinks moved from the original goal locations towards the new goals (Fig. 2g, h).

**Figure 2.**
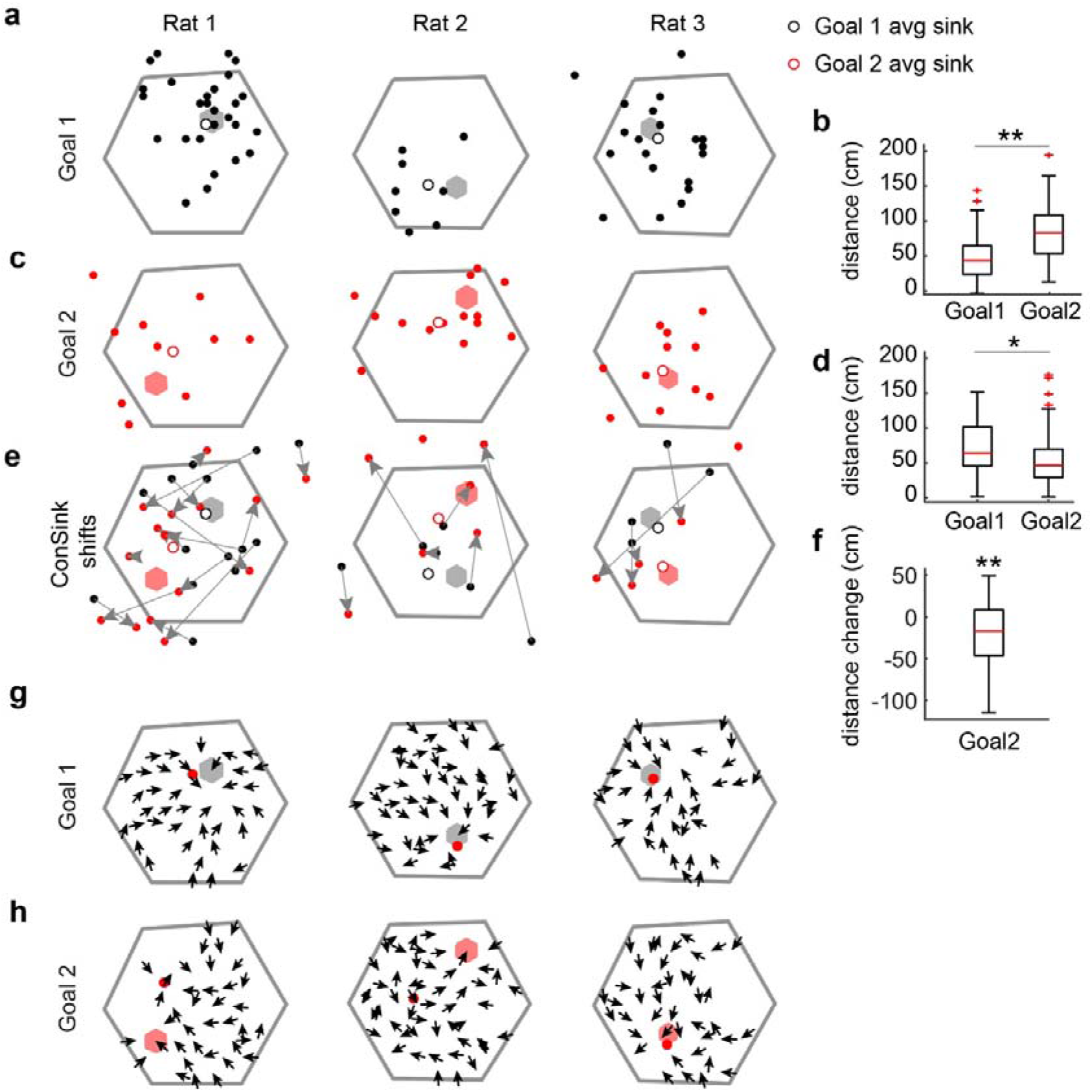
ConSinks are under the influence of goal location. **a**, Spatial distribution of ConSinks active only in Goal 1 (grey hexagon) before the goal switch. Average ConSinks, open circles. **b**, ConSink population was significantly closer to Goal 1 than to Goal 2 before goal switch (Wilcoxson rank sum test, p < 0.001). **c**, Spatial distribution of ConSinks active only in Goal 2 (red hexagon) after goal switch. **d**, ConSink population moved towards new goal (Goal 2) after goal switch (Wilcoxson rank sum test, p = 0.006). **e**, Arrows show movement of ConSinks from Goal 1 to Goal 2. **f**, ConSinks in (**e**) were closer to new goal after switch (Wilcoxson signed rank test, p < 0.001). **g**, Average vector fields for significant ConSink cells during Goal 1. **h**, as in **g**, but for Goal 2. Note that average ConSinks and vector fields shift towards the new goal.

Dividing the post-switch trials into 2 halves showed that ConSinks continued to cluster closer to the new goal with continued training (Fig. 3). Similarly, during continued training to the original goal on the first day of recording, they also moved closer to the goal (Extended Data Fig. 8). The distribution of ConSinks becomes more concentrated on the goal as the animal repeatedly navigates towards it, whether it is familiar or newly learned.

**Figure 3.**
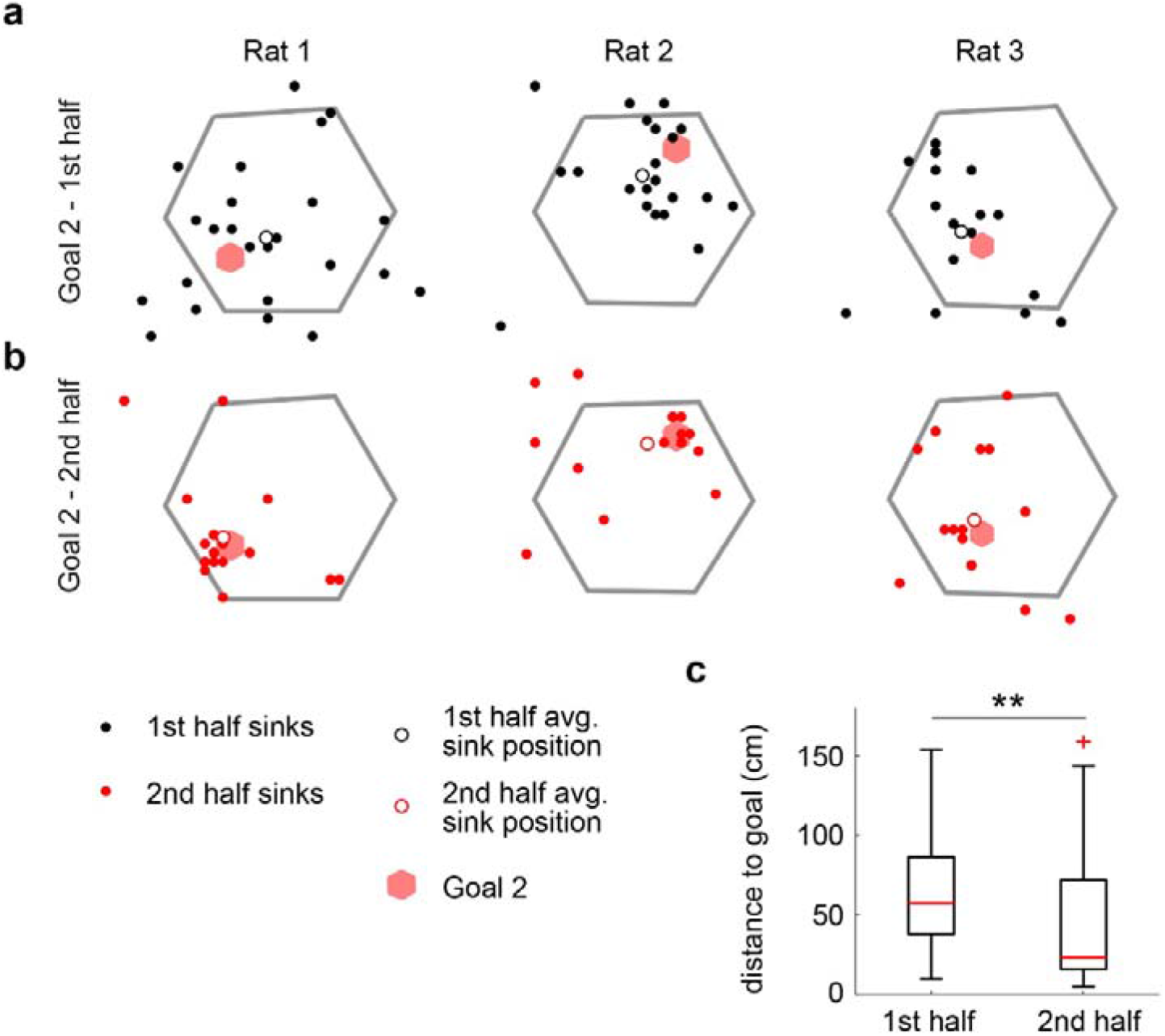
ConSinks continue to move closer to new goal during training. **a**-**c**, Spatial distribution of ConSinks during the first (**a**, Goal 2, 1^st^ half) and second (**b**, Goal 2, 2nd half) halves of Goal 2 training for each rat. Average ConSinks, open circles. **c**, ConSinks moved closer to the new goal in the second half of training (Wilcoxson sign rank test, p < 0.001).

The story was somewhat different for the place field centres; during navigation to the original goal they also clustered around it (Extended Data Fig. 9a-c), consistent with previous reports^9,13,14^. However, in contrast to the ConSinks, after the goal switch, place field centres did not reorganize around the new goal, although they shifted slightly in its direction. No such shift was seen in the place fields of non-ConSink cells (Extended Data Fig. 9d). We observed some remapping after the goal switch, but significantly less than that observed between the navigation task and open field foraging (see below, and Extended Data Fig.9e, f). In the honeycomb task, place field reorganization appears to be a slower process than that of ConSinks, perhaps reflecting different underlying processes.

## Place Cell ConSinks support navigation

We wondered whether the ConSink representation supported navigation on the honeycomb maze, and if so, how. Because the population vector fields show that firing is maximal when the animal is oriented towards the goal (see Fig. 1i), during unconstrained navigation, with the direct route to the goal available, the animal can simply follow the average population vector to the goal (direction G, Fig 4a thick red arrow). However, the fact that ConSink cells by definition have place fields suggests that they not only encode relative direction (Fig. 4b), but might also encode some combination of distance and allocentric direction to the sink (Fig. 4c-e); importantly, encoding of these extra variables could support calculation of the sink *positions* in addition to their relative *direction* from the animal and allow the calculation of the distance and direction vector to the goal. To test this, we used a previously published method (linear-nonlinear-Poisson nested models, abbreviated LN) that, from a pool of variables of interest, identifies those that significantly improve spike prediction from a fixed mean rate model^16^. We found significant encoding of various combinations of relative direction, distance, and allocentric direction to the sink in 70 of the 77 ConSink cells (Fig. 4f-j; Extended Data Fig. 10a). Of these cells, the largest group (28 of 77 cells) was comprised of cells significant for all 3 variables. 44 cells encoded a combination of variables sufficient for the calculation of sink positions (i.e. either all 3 variables, relative direction and distance, or allocentric direction and distance), while 26 cells encoded only relative direction, allocentric direction, or both. The lack of significance for any variables in the remaining cells (7 of 77) could be attributed to lower firing rates (Extended Data 10b). Each variable was well distributed across the population of ConSink place cells (Extended Data Fig. 10c-e), which therefore has all the information necessary to calculate the goal direction vector from any location. However, this simple strategy of following the goal direction vector is not adequate to solve the honeycomb maze task because the animal is frequently offered choices neither of which point directly to the goal. In such a case, the preferred choice that will take the animal closer to the goal is to select the path whose heading is most similar to the goal direction (the largest inner product). To identify this path from the alternatives, the animal needs to construct and compare the neural equivalents of the vector amplitudes in these directions (Fig. 4a, narrow red arrows). Although we have represented the mean allocentric direction of population spiking in the vector fields (see Fig. 1i), the underlying data can also be represented as a set of vectors with the average pointing towards the goal (Fig. 4k). We wondered whether this *vector set* provided enough information to allow the animal to choose between the different platforms. In general, the lengths of the vectors in the different directions decrease as an inverse function of the absolute angle from the average ConSink direction forming a *Fantail* configuration (Fig 4l). Choosing the direction with the largest population vector or its equivalent, the highest population firing rate, is the correct solution to the navigation problem.

**Figure 4.**
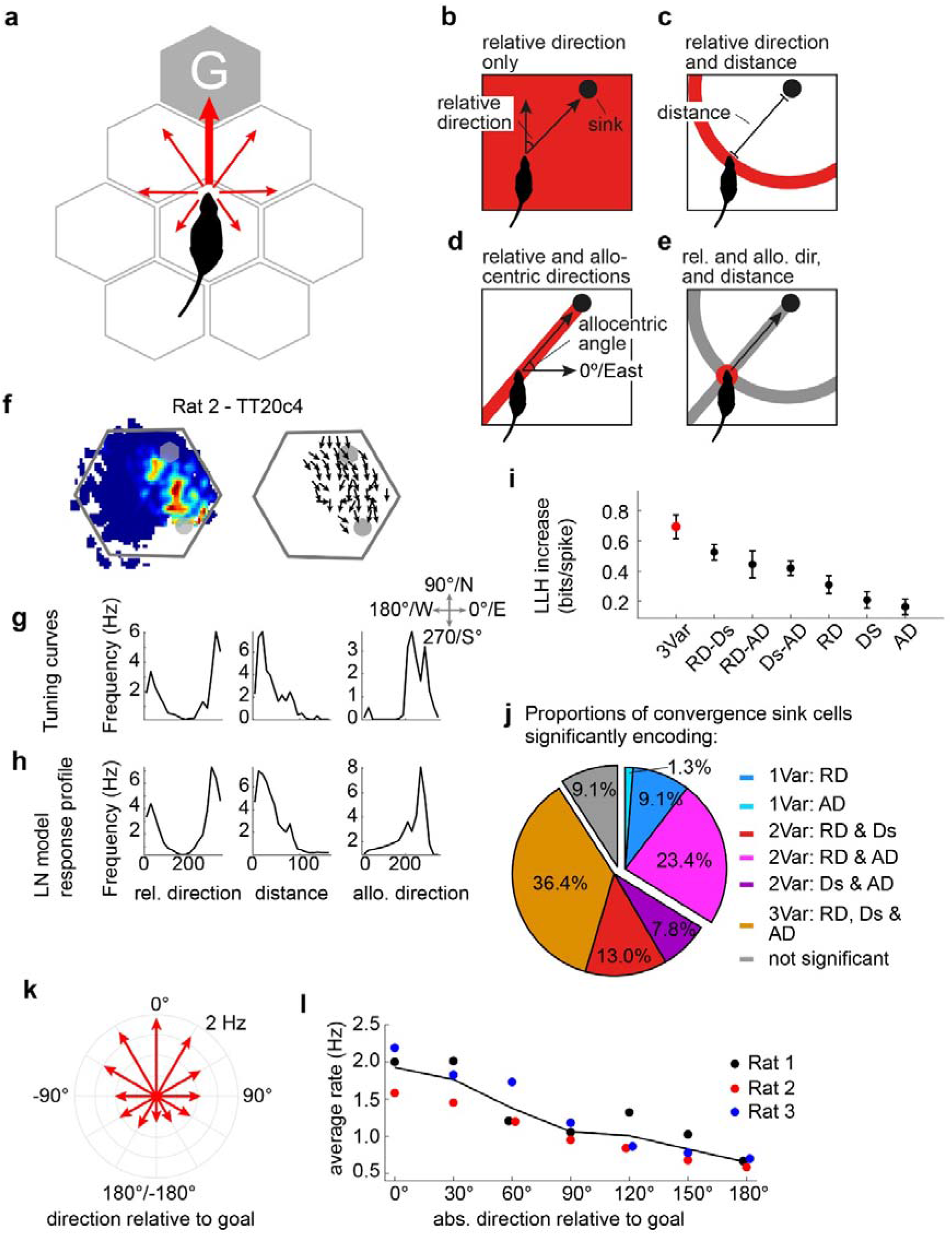
ConSink population firing patterns contain enough spatial information to solve the honeycomb maze navigation problem. **a**, Simple Fantail model predicts that firing rate vectors will be maximum in the direction of the goal and fall off monotonically with increasing angle from the goal. Information necessary to construct the goal direction vector consists of all or certain combinations of relative direction (**b**), distance (**c**), and absolute direction relative to the environment (**d**) to the ConSink, which together produce (**e**). **f**, Typical CA1 place cell with significant information coding for all three variables with raw tuning curves (**g**) and LN model response profiles (**h**). **i**, For the cell in (**f**), combination of all 3 types of information provides more information than any other combination (LLH, log-likelihood; 3Var, 3 variables; RD, relative direction; Ds, distance; AD, allocentric direction). Error bars = s.e.m. **j**, Percentages of ConSink cells encoding different combinations of the 3 variables in the LN model. **k**, Fantail data: population firing rate vectors across the 3 animals (red arrows) varying monotonically as a function of the angle between each platform direction and the population goal vector (Rayleigh test of non-uniformity of distribution, p < 0.001; mean direction = −1.50º). **l**, Population vectors for each animal conform to this model.

## Fewer ConSinks during open field foraging

Place cell firing is usually reported to be omnidirectional during random foraging in an open field with walls^6^ (although see^8^). During goal-directed behaviours on radial-arm mazes^8^ and linear tracks^16^, however, it is unidirectional, with cells firing in one direction and not the other, or in different places in the two directions. Omnidirectional pyramidal cells become unidirectional in the same environment when goals are introduced^17^. We recorded the same CA1 pyramidal cells from the single-goal honeycomb task during an open field foraging task on the same maze (all platforms in the raised position), and compared the place fields and ConSinks under the two conditions. 14% of the place cells (35/242) displayed significant tuning to convergence sinks in the foraging task, significantly fewer than during the navigation task (Extended Data Fig.11a-e). Only 6 cells had significant ConSinks in both tasks, and the shift in their sink locations, as well as preferred mean relative directions, across tasks indicated a complete reorganization of the hippocampal representation (Extended Data Figs. 11f-h). Unlike during navigation, the distribution of sinks was not denser around the navigation task goal location than expected by chance (Extended Data Fig. 11i). The existence of a single goal location in the navigation task as opposed to multiple random food sources in the foraging task is probably responsible for the remapping, although we cannot rule out the possibility that the difference might be due to the difference in the structures of the two maze configurations.

## Honeycomb maze navigation separates spatial representation from spatial action

Because the raising of the next choice platforms is delayed after each choice, the honeycomb maze presents an opportunity to observe the animal’s spatial representation before it chooses its next trajectory or even before it knows what choices will be offered. We examined 2 time periods: Wait Period 1, which started after the animal’s choice (see Fig. 1b) but *before* the next choice platforms had started rising (duration 4 sec); and Wait Period 2, which was chosen as the 4 second window leading up to the animal’s next choice (defined as the moment when the animal’s torso moves onto the chosen platform). During the wait periods, the rat systematically scans the environment, often turning through 360° with its head level with the horizontal plane (Fig. 5a, b), but frequently dipping down below the edge of the platform. We were particularly interested whether there might be differences in these representations between correct and error choices. We found that, on the former, the animal’s behaviour on average displayed even sampling of all directions relative to the goal, sometimes performing a complete 360° circuit. In contrast, prior to error choices, the distribution of behavioural orientations was heavily skewed away from the goal (Extended Figure 12). The firing rates of ConSink cells were significantly higher in correct choices both in the goal direction (Figure 5c, d), as well as in the preferred direction relative to their individual ConSinks (see Figure 1; Figure 5e). The corresponding Fantails were appropriately peaked towards the goal before correct choices during both periods but incorrectly rotated away from goal before incorrect ones. Thus, it appears that the fantail distributions relative to the goal predict subsequent behavioural choice even before any knowledge of the upcoming choices.

**Figure 5.**
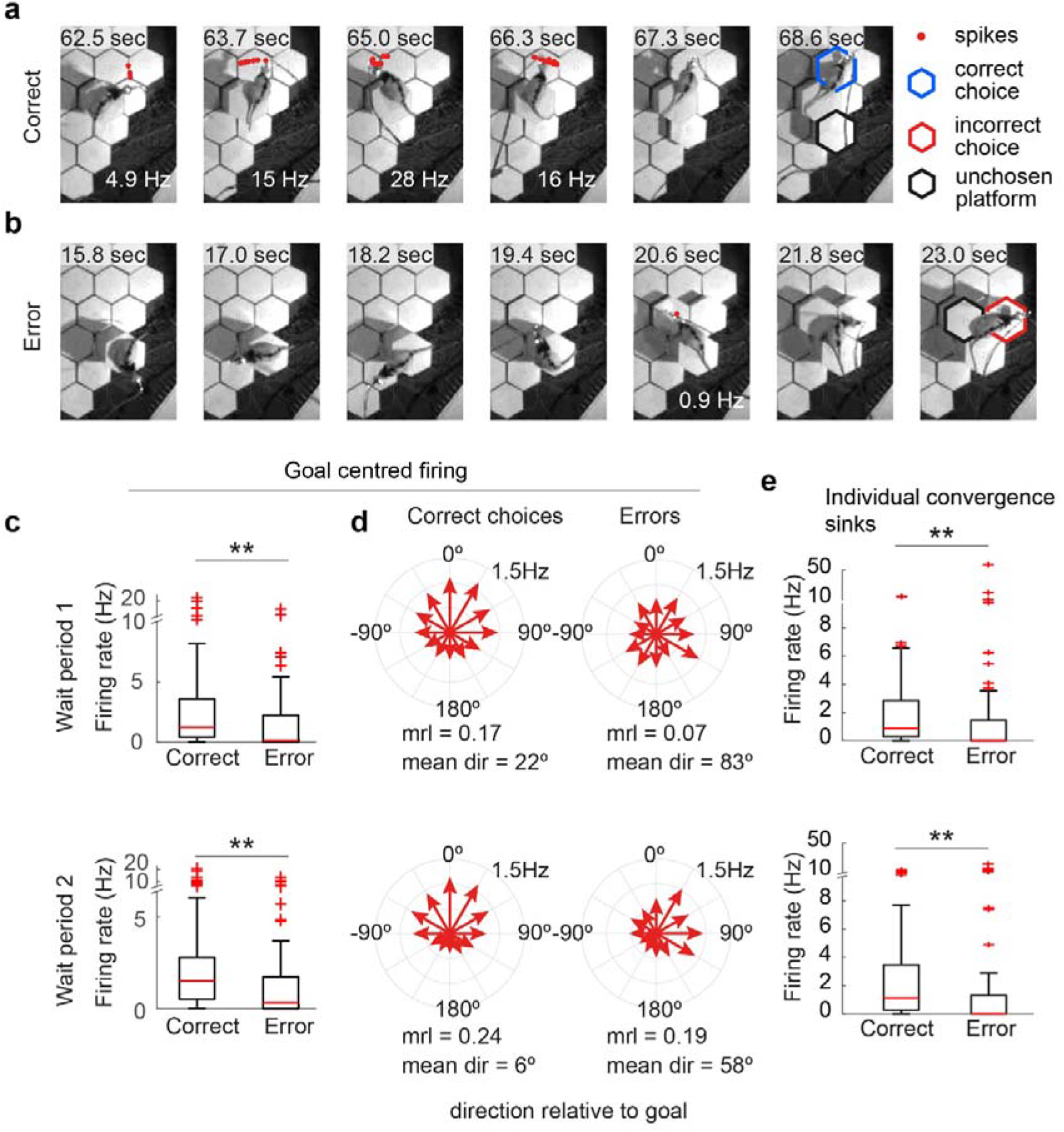
Goal-centred firing by ConSink cells is reduced on error sub-trials. **a**, Example ConSink cell (Rat 2, cell TT18c1, trial 16) immediately prior to correct choice. Note side-to-side scanning and robust firing in goalward direction before move onto the correct platform (goal beyond the top of the frame). **b**, Same cell as in (**a**) before incorrect choice. **c**, Firing rates in the goalward direction were reduced on error-trials, both during Wait Periods 1 (4 s period before raising the choice platforms; top) and 2 (4 s period before movement onto the chosen platform; bottom). Wilcoxson sign rank test, p < 0.001. **d**, Fantail plots of firing rates in directions relative to goal during correct choices have canonical forms, as in Fig. 4k, and peaks much closer to 0º than on incorrect choices. **e**, The tuning of ConSink cells to their individual sink positions is also disrupted on incorrect choices.

## Discussion

During navigation on the honeycomb maze, the firing patterns of a subset of CA1 hippocampal pyramidal cells are organized as vector fields oriented around a set of featureless environmental locations called ConSinks. While the total population of CA1 pyramidal cells provides information about the animal’s current location, the sub-population of ConSink cells contains all the information necessary for successful, flexible navigation in a familiar environment: allocentric information about the animal’s location in the environment (place coding), and both allocentric and egocentric information about distance, location and heading to the ConSinks, and, at a population level, Fantails describing the relative goalward value of different directions from any given location. While the ConSinks are arranged throughout the environment, they are concentrated around and centred near the goal, providing clear evidence for an effect of learning on ConSink location. This is further supported by the appearance of new sinks, and the disappearance or rearrangement of existing sinks, around a new goal after a goal-switch, as well as the continued movement of the sinks towards the goal during continued performance of the task within a single day. Finally, we find that firing towards the goal or the individual sinks is reduced and the fantail pattern altered on error trials, indicating that ConSink cells are crucially involved in navigation the goal.

We identify how the hippocampus can solve the honeycomb task using ConSink cells. When the direct route to the goal is available, because the ConSinks are densest around the goal, and, on average, the ConSink cells fire when their associated sinks are directly in front of the animal, the population firing rate is highest when the animal is oriented towards it. Thus the average vector field points to the goal and signals its distance. When the direct route to the goal is not available, the population rate falls monotonically with deviation from the direct goal heading, and the animal simply needs to move in the direction of highest firing rate afforded by the choices available, essentially comparing the lengths of the available branches of the Fantail.

On error choices, even before the new choice platforms rise, firing rates are reduced and the fantails are deviated from the canonical form. The pre-choice behaviour on these trials is also disrupted with the animal spending more time looking away from the goal than towards it as though its attention had been attracted elsewhere. They both recall the vicarious trial and error behaviour originally described by Meunzinger^18^ and Tolman^19^, and recently explored by Redish^20^ in the context of predictive place cell firing at the choice point. The Honeycomb maze reveals the 2D nature of the pre-choice ‘subjunctive’ representation.

Cells similar to the ConSink cells have been reported in mouse open field foraging^8^ and in humans performing a multiple object-in-location VR test^11^. In the present experiment, these occurred in only 14% of the cells during a foraging task, fewer than half of the percentage seen during the navigation task, and further, were not clustered around the concurrent navigation goal. While previous studies have reported the orientation of hippocampal formation cell firing relative to goals^10^, objects^21^, the centre of the environment^22^, or random points scattered around the environment^7^, the current work identifies a set of featureless locations dotted around the environment but organized around the goal, which they move closer to as learning proceeds. It must be left to future studies to determine how these Convergence Sinks are created and manipulated, and the elucidation of the properties of the underlying reinforcement mechanism. Examination of error choices suggests that selection of the correct choice platform cannot occur if the animal does not first activate a veridical representation of the hierarchy of choices and their valences. The process mediating the translation between spatial representation and spatial action must await further experimentation.

## Methods

### Subjects and surgical procedures

Subjects were 3 male Lister hooded rats purchased from Charles River Laboratories and aged between 9 and 12 months at the time of electrophysiological recordings. Animals were food deprived to ~85-90% of their baseline weight. All animal experiments were carried out in accordance with British Home Office Regulations (UK Animals Scientific Procedures Act 1986; Project License PPL PD8CBD97C to J. O’Keefe). Study protocols were in accordance with the terms of the Project License, which was reviewed by the Animal Welfare and Ethical Review Board at University College London.

### Surgery and recording

Rats were anesthetized with .5-1.5 % isoflurane. Craniotomies were made bilaterally over dHPC (4.2 mm posterior from Bregma, ±3 mm lateral from midline). The electrode array, containing 32 tetrodes (16 tetrodes per hemisphere), was implanted on to the surface of the cortex and electrodes turned 750 µm into the brain. One bone screw attached to the skull over frontal cortex served as ground and reference. Tetrodes (nichrome, ¼ Hard Pac coating, 0.0005 in diameter, Kanthal item #PF000591) were gold-plated to < 150 kΩ prior to implantation. Tetrodes were lowered to dorsal CA1 over two weeks, and rats continued to run daily training sessions on the track. Data was acquired using an Intan RHD USB interface board and RHD headstages.

### Maze

The honeycomb maze consists of 61 tessellated hexagonal platforms (11.5□cm each side) in an overall hexagonal configuration (total maze width ~200 cm). Each platform can be raised or lowered independently on a linear actuator, with the raised position ~30cm higher than the lowered position. Platforms were controlled with digital pulses generated in custom written software in LabView. The presence of an animal on a given platform was detected using load cells (RobotShop, cat. # RB-Phi-117) on which the platforms were affixed. The load cell signal was amplified with a custom made circuit and inputted into our custom LabView software.

The task was run as follows. First, a list of 13 start platforms was created in Matlab by randomly choosing a single platform from each of 13 maze subsections. The first start platform was then raised and the animal was manually placed atop it. The trial was then started from the custom LabView software by the experimenter. Two adjacent platforms were then pseudo-randomly selected by the software with 2 stipulations: first, that at least one of the platforms provided a position closer to the goal than the animal’s current position; and second, that previously unused platforms were selected from first, as long as the first stipulation could be met. The animal’s choice was registered once the load-cell system had registered its presence on one of the two choice platforms for a continuous 5 seconds. This triggered the lowering of the two other platforms, and after a delay of 4-10 seconds, another sub-trial commenced. In any sub-trial, the choice was registered as correct if the animal chose the platform closest to the goal. In some sub-trials, the two choice platforms were the same distance from the goal; these sub-trials were not included in the analysis of behavioural performance. Once the animal reached the goal platform, food reward (honey-flavoured corn flakes) in a small metal bowl was placed next to the animal. Once the animal had finished consuming the food reward, the experimenter placed the animal on a pedestal next to the maze. Every 4 trials, the maze was wiped down with 70% ethanol, and rotated 30 degrees (alternating between clockwise and counter-clockwise rotations) on a bushing located under the maze in order to prevent the animal from following scent trails to the goal.

The animal’s ability to correctly navigate the maze was confirmed using the binomial test^1^ to determine if the number of correct choices was greater than would be expected by chance given a 0.5 probability of correct choices.

In the first recording sessions (Fig. 1, Fig. 4, and Fig. 5), animals ran 13 trials of the task, followed by open field foraging, and then a second set of task trials that varied in number between the 3 animals (13 additional trials for Rat 1, 7 additional trials for Rat 2 and 3). In the second recording sessions. In the second recording sessions, (Fig. 2 and Fig. 3), animals ran 13 trials to Goal 1, followed by 13 trials to Goal 2; there was a set of “learning” trials interleaved between the two sets, detailed below in the section entitled “*Goal switch training*”.

### Spike sorting

Spikes were automatically sorted using KiloSort^2^, followed by manual refinement using Phy^3^ which consisted mainly of merging and deleting clusters, using autocorrelations and cross-correlations as a guide. Cells with greater than 1% of spikes within the first 2 ms of the spike autorcorrelation were excluded from further analysis. Cells were classified as pyramidals, interneurons, or left unclassified (excluded from further analysis) on the basis of spike width, mean rate, burst index (nSpikes from 0 – 10 ms of the autocorrelation / nSpikes from 40 – 50 ms of the autocorrelation), and oscillation score (Muresan et al. 2008) using a PCA analysis. Principal components were calculated from the 4 variables, and the first 2 principal components plotted as a scatter plot. Pyramidal cells tend to cluster together, while interneurons are scattered outside the main cluster; the experimenter selected the cells within the cluster by manually drawing a boundary, followed by visual verification of the waveforms.

### Video tracking

Video was recorded in custom LabView software using a monochrome usb camera at a frame rate of ~25 frames per second (Imaging Source). Tracking was performed offline using DeepLabCut^4^. In DeepLabCut, we trained the network to identify 2 infrared LEDs positioned on top of the animal’s implant, as well as dark fur patches on the shoulders and back. The animal’s head position was taken as the midpoint between the 2 LEDs, and his head angle was taken as the angle of the line between the LEDs.

### Histology

Marking lesions were made using 20 µA of anodal current for 10 seconds. Animals were transcardially perfused with phosphate-buffered saline followed by 4% paraformaldehyde (PFA), brains cryo-protected in 30% sucrose/4% PFA, and frozen slices of 40µm cut. Slices were stained with Cresyl Violet.

### Pre-processing spike data

Hippocampal place cell data is typically velocity-thresholded since place cells lose some place tuning when the animal stops moving. In our task, because the animal was not able to move freely around the maze, this seemed an inappropriate approach. Instead, we focussed on excluding sharp wave ripple events by using 3 criteria that had to be met simultaneously: 1) theta power (6-12 Hz) below the mean; 2) population firing rate 2 standard deviations above the mean; 3) ripple power (100-250 Hz) 2 standard deviations above the mean. Spectral power was computed by taking the absolute value of the output of the continuous 1-D wavelet transform (Matlab function ‘cwt’). Data were only excluded if all 3 criteria were met for a minimum duration of 50 ms. Spike data from a given cell was only used for analysis during any of the conditions (honeycomb task, forage, goal1, goal2) if it fired at least 500 spikes in the relevant condition after this exclusion.

### Relative direction analysis

The field of view was tiled with potential convergence sinks, arranged along the x and y axes at ~7 cm intervals (34 × 29 total positions). The head directions relative to each potential sink were then calculated for each spike by subtracting the angle of the vector from the animal’s position to the sink position from the animal’s allocentric head direction. Thus, if the animal was facing the sink, these two directions were equal, and the relative direction was 0°. If the animal was facing in the opposite direction, the relative direction was 180°. The convention used in this paper is that positive relative directions indicate that the animal’s head direction was to the left a line from the animal to the sink (i.e. the sink is to the animal’s right), and negative directions indicate a rightward head direction relative to the sink. For a given cell, a binned distribution of relative directions could then be calculated (24 bins spanning −180° to 180°).

This distribution then had to be corrected for uneven sampling of relative direction by the animal. Because of the potential for differences in sampling of relative direction across the maze, we calculated control distributions of relative direction at each platform (61 platforms total) using all video frames in which the animal occupied a given platform (an animal was deemed to be occupying a platform if his torso was within the platform’s perimeter). For each cell, the distributions were summed according to the number of spikes the cell fired on each platform. Finally, the cell’s relative direction distribution could then be divided by the summed control distribution, providing a corrected distribution taking into account any uneven sampling of relative direction by the animal across the spatial extent of the maze. From this corrected distribution, using the CircStat toolbox^5^, we computed the Rayleigh test for non-uniformity of circular data (all ConSink cells were significantly non-uniform) and calculated the mean direction and the mean resultant length (MRL). Thus each cell had an MRL value associated with each potential sink; the candidate sink was taken as the potential sink with the highest MRL.

To test whether a cell was significantly tuned to direction relative to the candidate sink, we used a bootstrap method in which we shuffled the cell’s head directions such that the head directions were no longer associated with their actual positions on the maze. Distributions were calculated as above, yielding MRL values for each xy position. For each of the 1000 shuffles, the maximal mrl value across all xy positions was used to make a distribution of MRL values. A cell was deemed to be significantly modulated by relative direction if its MRL was greater than the 95^th^ percentile of the control distribution (see Extended Data Fig. 4).

Because place cells frequently fired in bursts as the animal scanned the environment (e.g. see Extended Data Fig. 2), causing a smearing of head direction that could potentially lead to false negatives in our search for ConSink cells. Thus, we repeated our search for ConSink cells using only bursts, and averaging relative direction and position within each burst to eliminate the smearing effect. Bursts were defined as trains of at least 10 spikes fired with interspike intervals less than or equal to 0.25 sec. If two bursts were separated in time by less than 0.5 sec, they were combined. If a cell was significant in both analyses (15 of 77 cells), only the data from the burst analysis, which produced the greatest tuning in all cases, was carried forward into subsequent analyses. In these 15 cells, we confirmed that both analyses identified the same ConSink positions (distance between ConSinks, within same cells = 10.3 cm, different cells = 85.2 cm, p < 0.001; median difference in preferred relative direction, within same cells = 3.3º, different cell = 74.9º, p < 0.001).

To make the vector fields for individual ConSink cells (e.g. Fig. 1d-g), the field of view was binned into 20 × 16 spatial bins, and a mean head direction value was calculated for each bin with greater than 20 spikes. For population vector fields, bins instead corresponded to maze platforms.

### Behavioural analysis

To determine what platform an animal was on for *post-hoc* analysis, we tracked the position of the animal’s torso using a dark fur patch behind the shoulders in DeepLabCut^4^. The animal was considered to be on a particular platform if the torso position was within the platform’s perimeter. For the analysis of correct and error choices (Fig. 5), Wait Period 2 was defined in relation to the time when the animal’s torso moved onto the chosen platform, and was taken as the 4 sec window starting 5 seconds before this transition; the 1 second gap between the end of Wait Period 2 and the transition to the new platform ensured no contamination of Wait Period 2 by the transition itself. Wait Period 1 was defined as the time after the previous sub-trial start and unchosen platforms had lowered and before the next choice platforms were raised.

### Goal switch training

In the goal switch trials, all animals ran 13 trials to Goal 1. Once completed, it was necessary to teach the animals that the goal position had switched. To do this, we then ran a number of “easy” trials to Goal 2. These easy trials were characterized by choice platforms that all led the animal closer to the new goal, such that the animal would arrive at the new goal through no real choice of his own. Once he arrived at the new goal, he was rewarded as normal. These trials were interleaved with easy unrewarded trials to Goal 1.

The training sequences for each animal were as follows. Rat 1 ran 4 easy trials to Goal 2, followed by a normal trial which he was not able to complete successfully. He then ran 2 easy unrewarded trials to Goal 1, followed by a single easy rewarded trial to Goal 2. He subsequently ran 13 normal trials to Goal 2, all of which are included in the presented analysis.

The data presented for Rat 2 is his second goal switch session. His first goal switch session was not completed successfully due simply to his inability to learn the new goal location. We subsequently ran brief sessions across 6 days of 1-4 trials to this new goal location in which he demonstrated clear learning of this new goal location. We then ran a second goal switch session, using Goal 2 from the failed goal switch session as Goal 1. After running 13 trials to Goal 1, we switched the goal and the animal ran 3 easy rewarded trials. He subsequently ran 15 normal trials to Goal 2. The final 13 of these trials were used in all analyses except for the analysis examining movement of the sinks across the two halves of the Goal 2 epoch (Fig. 3), which used the first 13 trials.

Rat 3 ran 13 trials to Goal 1, followed by 8 easy trials alternating between rewarded trials to Goal 2 and unrewarded trials to Goal 1. This was followed by 17 normal trials to Goal 2. The final 13 of these trials were used in all analyses except for the analysis examining movement of the sinks across the two halves of the Goal 2 epoch (Fig. 3), which used the first 13 trials.

### LN analysis

To determine the dependence of ConSink cell spiking on distance and direction to the sink, we used a technique developed to identify mixed-selectivity in individual neurons by quantifying the dependence of spiking on all possible combinations of a set variables^6^. A model, corresponding to a particular combination of variables, is trained by optimizing a set of parameters that convert animal state vectors corresponding to the variables of interest into firing rates, which are estimated as an exponential function of the sum of each variable value projected onto the set of parameters. The analysis uses 10-fold cross-validation, splitting the data into 10 equal sized partitions, training the model using 9 of the partitions, and testing on the held out partition such that each partition is tested once. The log-likelihood (LLH) increase in spike prediction relative to a mean firing rate model is calculated for each model, and the simplest model (i.e. fewest number of variables) that produces a significant increase relative to the mean firing rate model, as well as, in the case of multi-variable models, a significant increase over any simpler models (i.e. models with fewer variables) is selected as the model that best describes the neuron’s tuning. The significance is assessed using Wilcoxon signed rank tests comparing the LLH increases for each test partition across the relevant models. We adapted the model to use 3 variables: 1) relative direction to the sink (RD), 2) distance to the sink (Ds), 3) direction from animal’s position to the sink (AD); this produced 7 possible models (i.e. a 3-variable model, 3 2-variable models, and 3 single-variable models). Firing rate and animal state vectors were constructed using 100 ms windows. Relative direction and direction from position were both binned using 18 bins spanning −180° to 180° and 0° to 360°, respectively. Distance to sink was binned using 20 bins from 0 cm to the animal’s maximum distance to the sink.

### Fantail plots

To calculate the fantail plot (Fig. 4k) showing the population firing rates in the range of head directions relative to the goal, spikes within animals were combined across all ConSink cells. For each spike, the animal’s head direction relative to the goal was calculated in the same way as for head direction relative to the ConSink (see *Relative direction analysis* above). Spikes were then separated according to the platform occupied by the animal during spiking, and for each platform, its associated spikes were binned according to relative direction to goal (30º wide bins). Similarly, for each platform, the total amount of time that the animal spent within each relative direction bin was determined. Finally, the spike counts in each bin were divided by the total time (in seconds) spent in each bin, to produce firing rates in each bin. These were then averaged across all platforms and divided by the total number of ConSink cells to generate a per cell firing rate in binned direction relative to the goal; these values are shown in Fig. 4i, and are averaged across animals in Fig. 4k.

### Remapping analysis

To assess remapping between the honeycomb task and open field foraging, we created rate maps for all cells by partitioning the field of view into 1280 bins (40 bins in x direction, 32 bins in y direction). To establish baselines for cell stability, to which we could compare our remapping data to assess significance, we created rate maps corresponding to the first and second halves of the task and open field foraging epochs; specifically, for each spatial bin, we calculated the total occupancy (in seconds) and placed the data corresponding to the first and second halves in the corresponding rate maps. Population vector correlations were performed by constructing vectors for each bin using the firing rates of each cell at that bin and then calculating Pearson’s linear correlation coefficient for the 2 vectors being compared. Similarly, place field correlations were performed by linearizing the 2 dimensional rate maps for a given cell, and calculating the correlation between the 2 vectors.

### Place field centres

Place fields centres were taken as the centre of mass of the cell’s rate map.

### Statistical procedures

All statistical tests were two-sided and non-parametric unless stated otherwise. In box plots, the central mark indicates the median, and the bottom and top edges of the box indicate the 25th and 75th percentiles, respectively. The whiskers extend to the most extreme data points within 1.5 times the interquartile range away from the bottom or top of the box, and all more extreme points are plotted individually using the ‘+’ symbol.

## Acknowledgements

We thank Del Halpin for building the honeycomb maze and, with Robb Barrett, for help prototyping the platforms and modifying the maze; Piotr Sienkiewicz for designing and building the electronics; Graeme McPhillips and Martyn Stopps for consulting on the load cell circuitry; Aleks Radulovic for help with data collection; Borgel Greenaway and Jessica Broni-Tabi for histology; and Marius Bauza and members of the O’Keefe lab for helpful discussions regarding the results and manuscript. This work was supported by the Sainsbury Wellcome Centre Core Grant from the Gatsby Charitable Foundation and Wellcome Trust (090843/F/09/Z), and a Wellcome Trust Principal Research Fellowship (Wt203020/z/16/z) to JOK.

## Author Contributions

J. O. and J. O. K. designed the experiments. J. O. collected the data. J. O. and J. O. K. analysed the data and wrote the manuscript.

## Competing interests statement

The authors declare no competing interests.

## Additional Information

**Supplementary Information** is available for this paper.

## Notes

### Competing Interest Statement

The authors have declared no competing interest.

