## Supplemental Data for "Hippocampal place cells use vector computations to navigate"

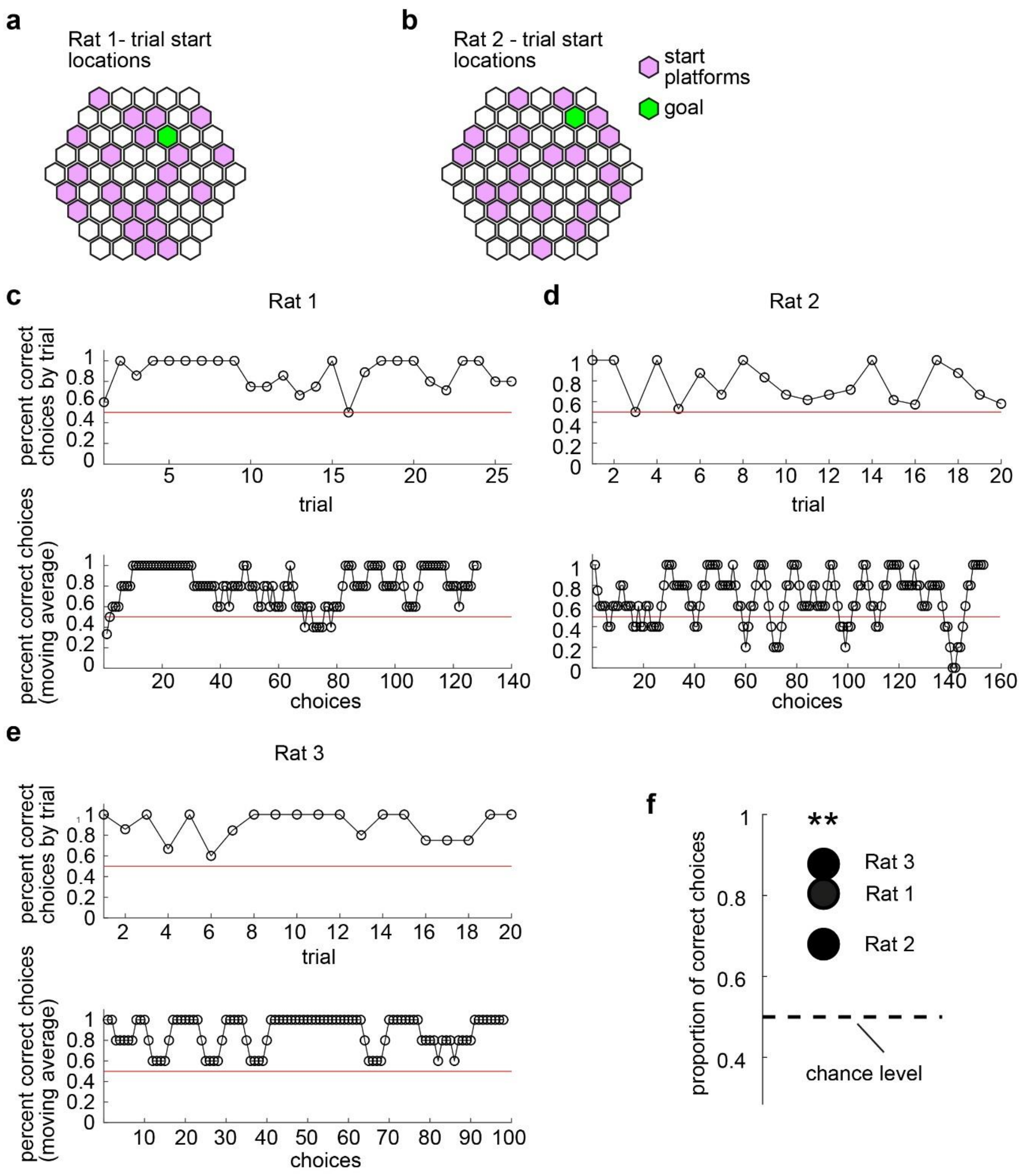


**Extended Data Figure 1 │ Behavioural Summary. a**, Schematic of the honeycomb maze showing all start platforms and the goal platform from Rat 1’s session. b, Same as is in (**a**) but for Rat 2. **c**, Top panel, Percentage of choices that were correct averaged by trial for Rat 1. Bottom panel, Running average of correct choices (every 5 consecutive choices). **d**, **e**, As in (**c**) but for Rats 2 and 3, respectively. **f**, The total proportion of correct choices for each rat. Each rat made significantly more correct than incorrect choices; $p < 0.001$, binomial test within each animal.

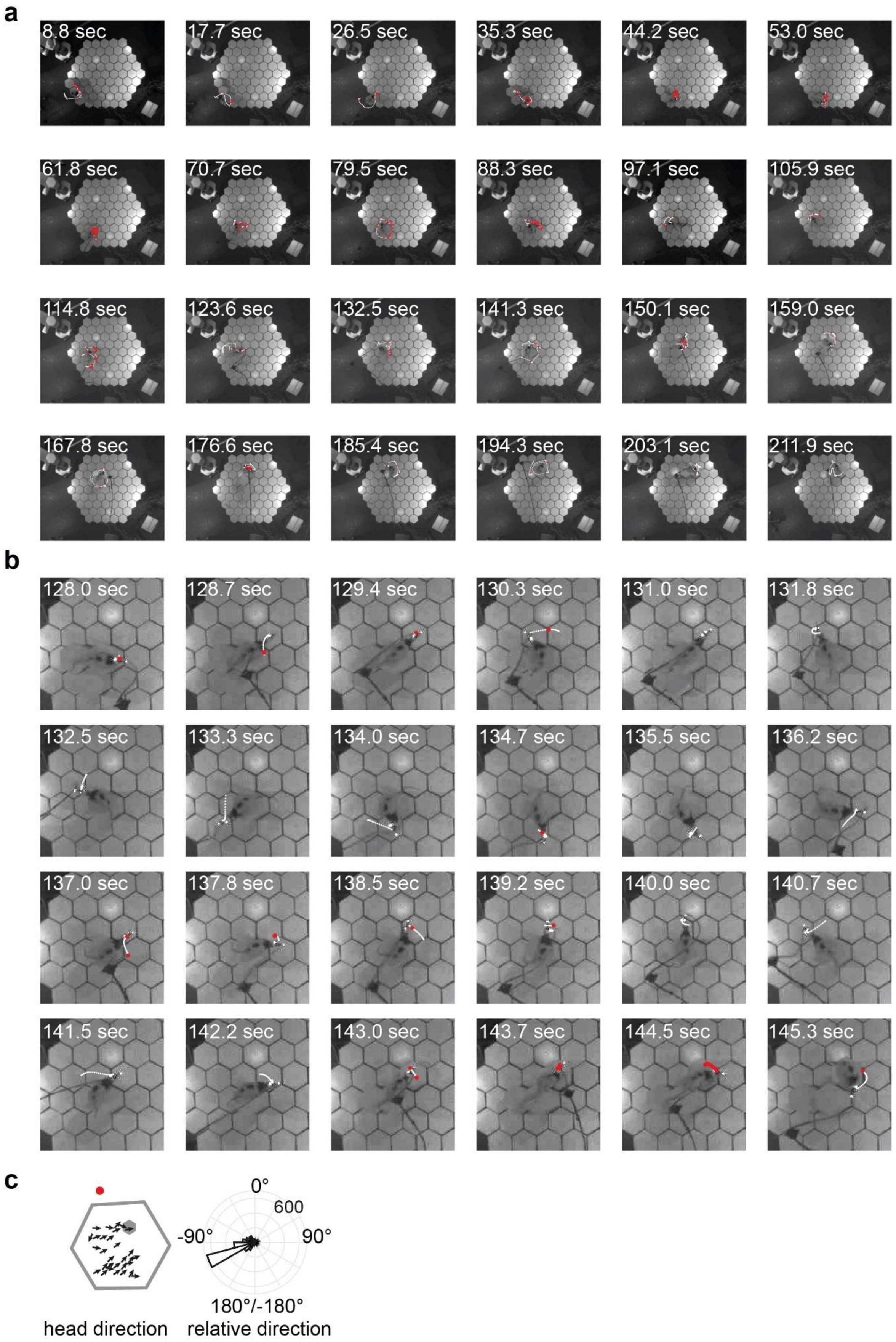
a
8.8 sec
17.7 sec
26.5 sec
35.3 sec
44.2 sec
53.0 sec
61.8 sec
70.7 sec
79.5 sec
88.3 sec
97.1 sec
105.9 sec
114.8 sec
123.6 sec
132.5 sec
141.3 sec
150.1 sec
159.0 sec
167.8 sec
176.6 sec
185.4 sec
194.3 sec
203.1 sec
211.9 sec
b
128.0 sec
128.7 sec
129.4 sec
130.3 sec
131.0 sec
131.8 sec
132.5 sec
133.3 sec
134.0 sec
134.7 sec
135.5 sec
136.2 sec
137.0 sec
137.8 sec
138.5 sec
139.2 sec
140.0 sec
140.7 sec
141.5 sec
142.2 sec
143.0 sec
143.7 sec
144.5 sec
145.3 sec
c
0°
600
-90°
90°
180°/-180°
head direction
relative direction

**Extended Data Figure 2 │ Typical behaviour and spiking of a ConSink neuron.**

**a**, Screengrabs of a complete trial of the honeycomb task (Rat 1, trial 3). In white is the path of the animal from the time of the previous screengrab (or from 0 sec for the first screengrab) to the time of the current screengrab. In red are the spikes fired by a representative ConSink cell (TT15c6). **b**, Screengrabs from a portion of the same trial in (**a**) at a higher temporal resolution. Note how the animal completes a full 360° rotation while waiting for the next pair of platforms to be presented (platforms start to raise at 142sec); this is typical of all 3 animals' behaviour. **c**, The same cell's vector field (left) and the polar plot of spike directions relative to the convergence sink (right; the sink is plotted in red at left).

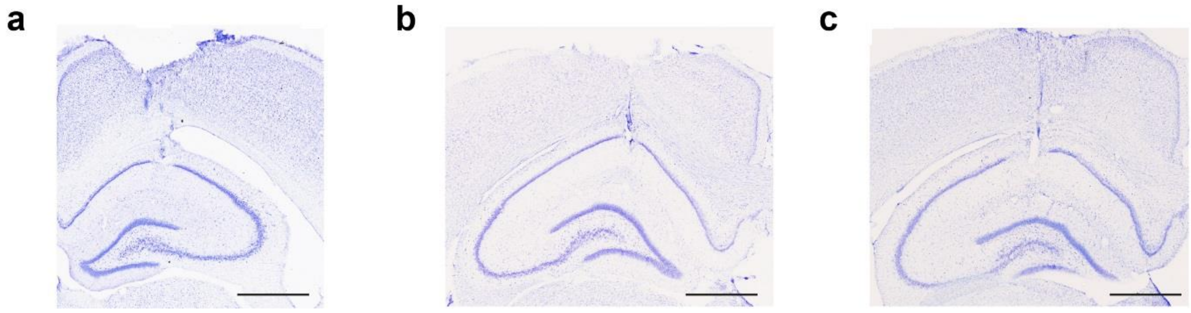


**Extended Data Figure 3 │ Histology showing tetrode position in the CA1 cell layer in dorsal hippocampus.** Example cresyl violet stained coronal slices showing tetrode marking lesions in Rat 1 (**a**), 2 (**b**), and 3 (**c**). Scale bar = 1000 µm.

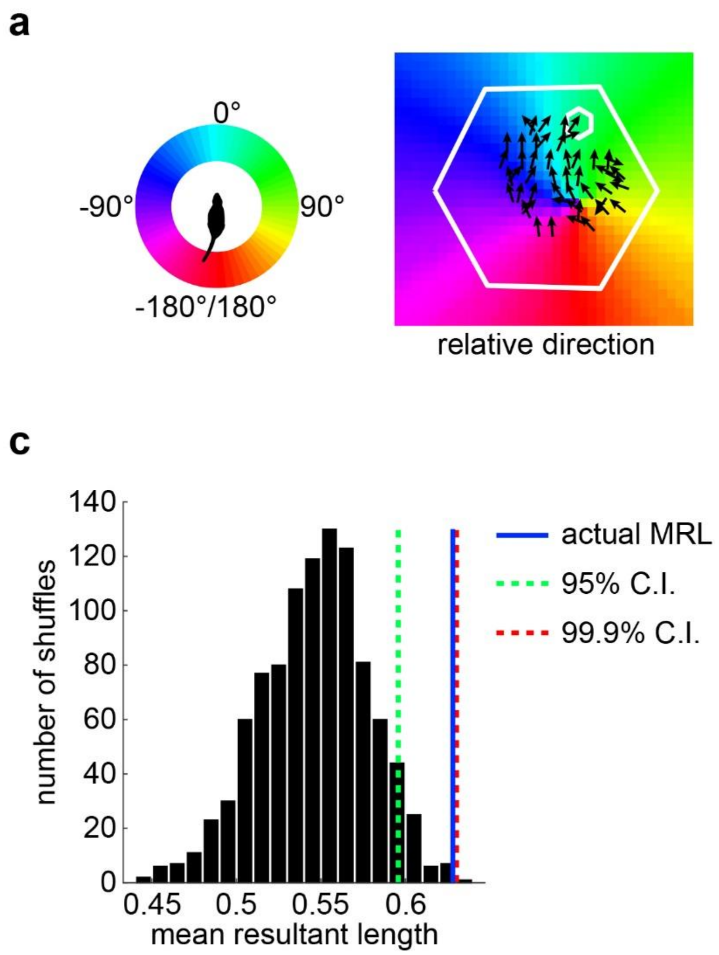


**Extended Data Figure 4 │ Calculation of the ConSink. a**, Left, Each division of the colour wheel represents 1 search position in a polar co-ordinate framework centred on the animal's head with 0° straight ahead. Right, The spatial environment is tiled by candidate sink positions. At each candidate position, the direction of all spikes relative to that position can be calculated (by subtracting the direction of the vector from the animal to the candidate position from the animal's allocentric head direction). From this, a mean direction can be calculated, and is plotted here according to the colour wheel at left. The vector field (i.e. the mean allocentric direction of spikes at binned spatial positions) of an example ConSink cell (Rat 2, cell TT18c1) is overlaid. **b**, At each candidate sink position, the mean resultant length (MRL) of the associated distribution of relative directions is calculated (candidate positions are colour coded by MRL value). The candidate position with the largest MRL, which indicates the concentration of the polar distribution in the mean direction for that position, is taken as the ConSink (black closed circle; MRL = 0.63). **c**, To determine whether a cell was significantly modulated by relative direction to the candidate position identified as in (**b**), we shuffled the allocentric head directions associated with each spike, and recalculated relative direction MRLs at each search position, using the maximal MRL for our control distribution. This procedure was repeated 1000 times, and confidence intervals constructed. A cell was deemed to be significantly modulated if it's MRL was greater than the 95$^{th}$ percentile of the shuffled distribution of maximal MRLs.

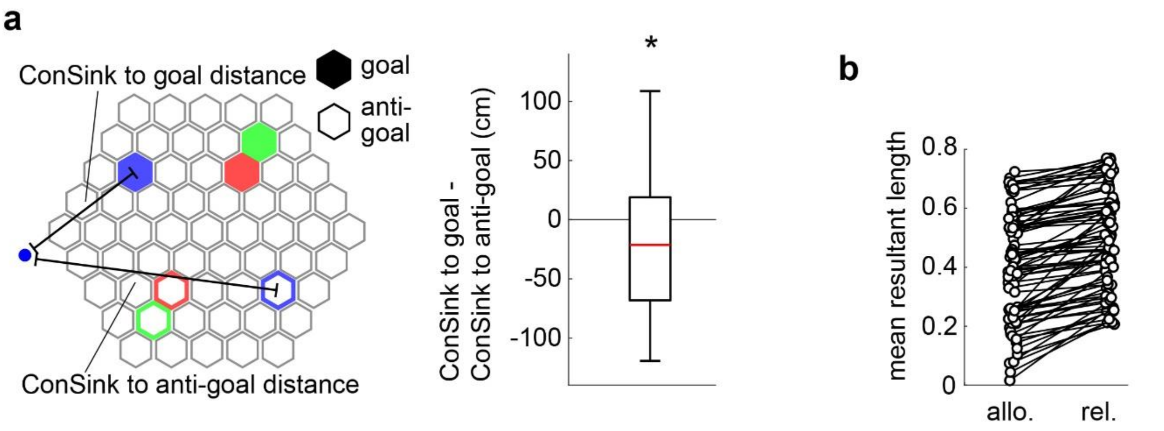


**Extended Data Figure 5 │ ConSinks cluster around the goal. a**, Left, Schematic of the maze showing positions of goals (red: Rat 1; green: Rat 2; blue: Rat 3) and platforms used as anti-goals in the analysis at Right. The anti-goal positions were produced by mirroring the goal positions across the axis perpendicular to a line between the maze vertex closest to the goal and the opposite vertex. For each relative direction cell, the distances from its convergence sink to the goal and anti-goal were calculated. The differences between these distances is plotted at Right. Convergence sinks were closer to the goal than the anti-goal, suggesting greater density around the goal (Wilcoxon signed rank test, $p = 0.0147$). **b**, MRLs calculated from allocentric (allo.) and relative (rel.) head directions for each relative direction cell during task. Note that for every relative direction cell, the MRL calculated from relative directions is greater than the MRL calculated from allocentric head directions.

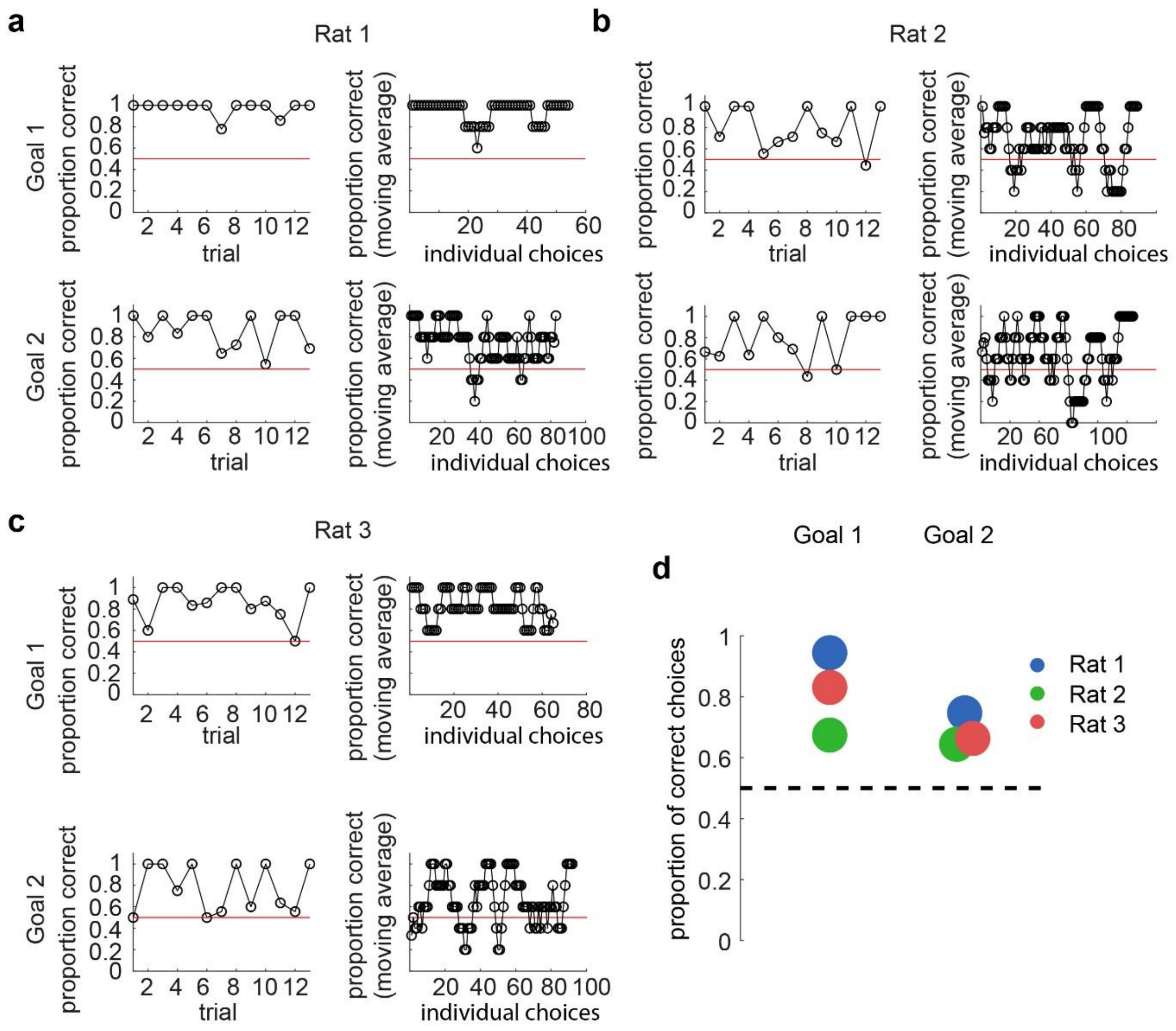


**Extended Data Figure 6 │ Animals successfully navigated to new goal locations in the goal switch sessions. a**, Top left, Proportion of choices that were correct to Goal 1 averaged by trial for rat 1. Top right, Running average of correct choices (averages of every 5 consecutive choices) to Goal 1 for rat 1. Bottom right and left, as at top, but to Goal 2. **b**, **c**, As in (**a**) for rat 2 and rat 3, respectively. **d**, The total proportion of correct choices for each rat for goals 1 and 2. Each rat made significantly more correct than incorrect choices for both goals; Rat 1, goal 1: $p < 0.001$, goal 2: $p < 0.001$; Rat 2, goal 1: $p = 0.0013$, goal 2: $p = 0.0042$; Rat 3, goal 1: $p < 0.001$, goal 2: $p = 0.0023$; binomial test within each animal.

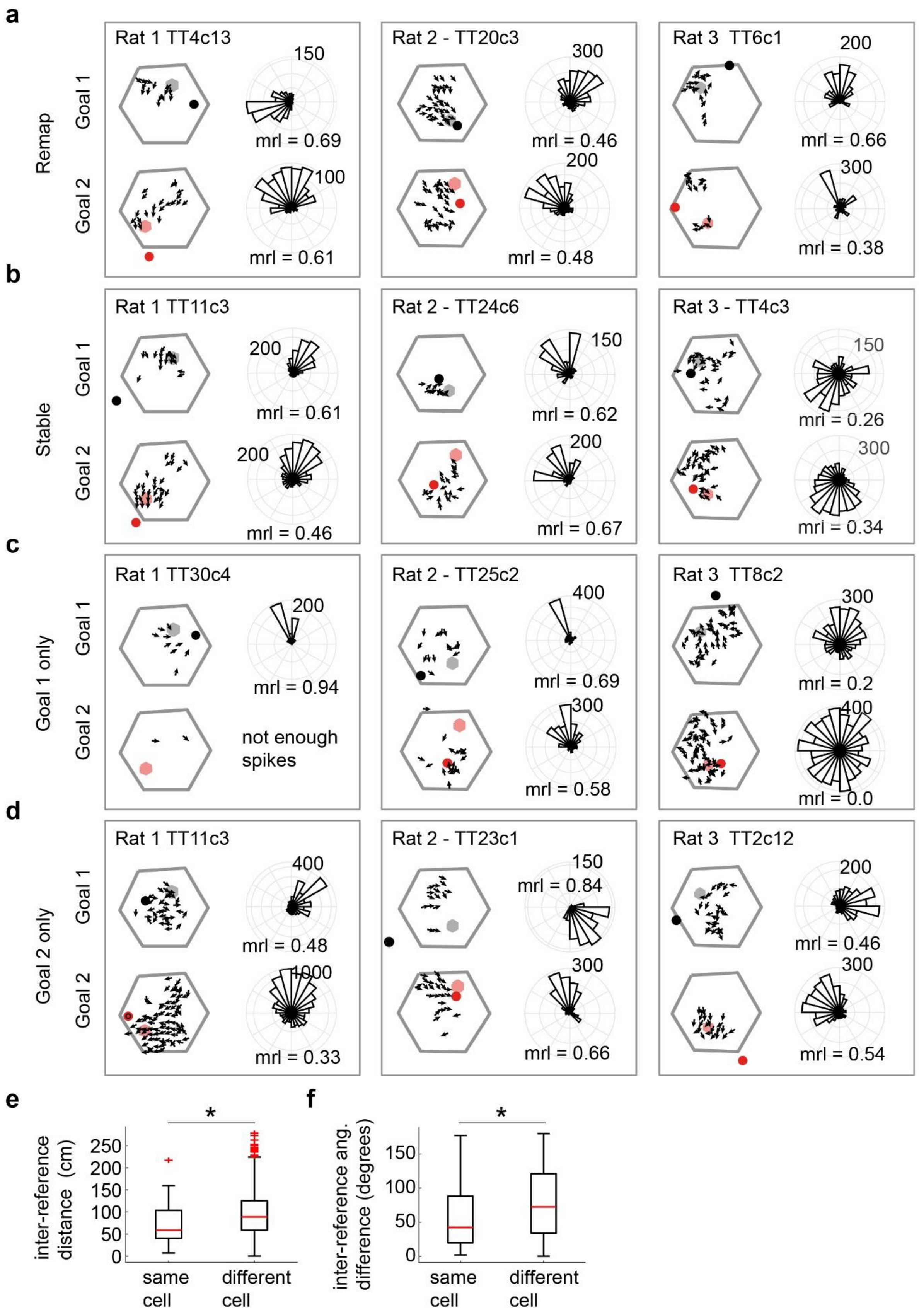

a
Remap
Rat 1 TT4c13
Rat 2 - TT20c3
Rat 3 TT6c1
Goal 1
Goal 2
mrl = 0.69
mrl = 0.61
mrl = 0.46
mrl = 0.48
mrl = 0.66
mrl = 0.38
b
Stable
Rat 1 TT11c3
Rat 2 - TT24c6
Rat 3 - TT4c3
mrl = 0.61
mrl = 0.46
mrl = 0.62
mrl = 0.67
mrl = 0.26
mrl = 0.34
c
Goal 1 only
Rat 1 TT30c4
Rat 2 - TT25c2
Rat 3 TT8c2
mrl = 0.94
not enough spikes
mrl = 0.69
mrl = 0.58
mrl = 0.2
mrl = 0.0
d
Goal 2 only
Rat 1 TT11c3
Rat 2 - TT23c1
Rat 3 TT2c12
mrl = 0.48
mrl = 0.33
mrl = 0.84
mrl = 0.66
mrl = 0.46
mrl = 0.54
e
inter-reference distance (cm)
same cell
different cell
f
inter-reference ang. difference (degrees)
same cell
different cell

**Extended Data Figure 7 │ Examples of ConSink cells recorded during the goal switch sessions. a**, Example cells that were significantly modulated during both Goal 1 and Goal 2 epochs, but whose ConSink changed position. Vector fields (left) depict mean head direction at binned spatial locations. The ConSink and goal location depicted in black (Goal 1) or red (Goal 2). Right, Polar plot showing the distribution of head directions relative to the ConSink. **b**, As in (**a**), but examples whose ConSink positions did not change. **c**, Examples of cells that were only significantly tuned during Goal 1. Where sufficient spikes were fired (min. 500), the best candidate ConSink position for Goal 2 is plotted. **d**, As in (**c**), but examples that were only tuned during Goal 2. **e**, Distances between Goal 1 and Goal 2 sinks within cells that had sinks during both epochs (e.g. the cells shown in (**a**) and (**b**)) were smaller than those between all other possible pairs of Goal 1 and Goal 2 ConSink cells, indicating stability of sink position within this sub-population across the goal switch. Wicoxon rank sum test, $p = 0.019$. **f**, Differences in mean direction between Goal 1 and Goal 2 sinks within cells that had sinks during both epochs (e.g. the cells shown in (**a**) and (**b**)) were smaller than those between all other possible pairs of Goal 1 and Goal 2 ConSink cells, indicating stability relative direction within this sub-population across the goal switch. Wilcoxon rank sum test, $p = 0.041$.

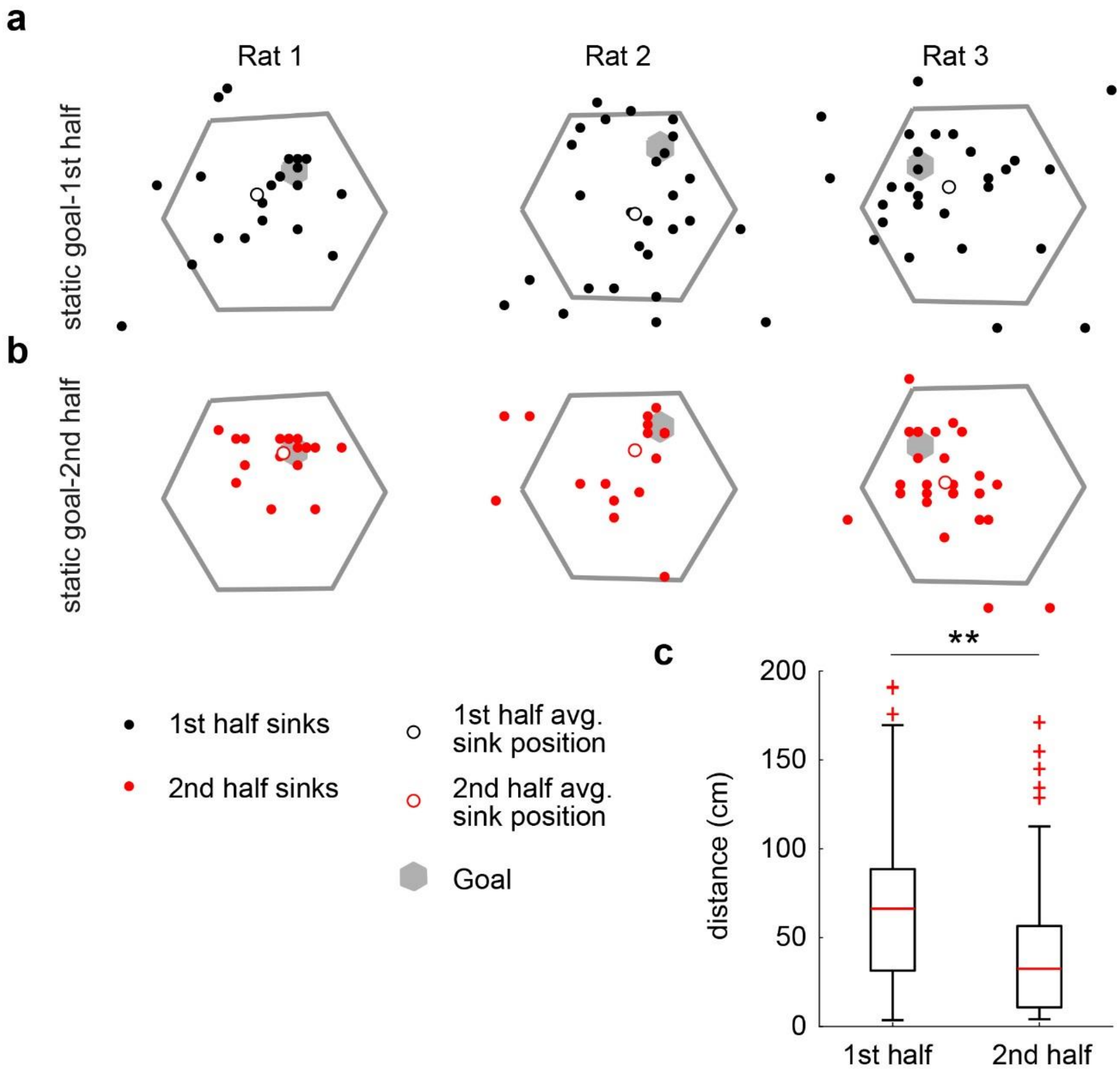


**Extended Data Figure 8 │ ConSinks move closer to the original goal with experience within a day.** Compared to ConSinks calculated during the first half of the first recording session (**a**), ConSinks calculated during the second half of the first session (**b**) appear more concentrated around the goal. **c,** The second half ConSinks are significantly closer to the goal than first half ConSinks (Wilcoxon signed rank test, $p < 0.001$).

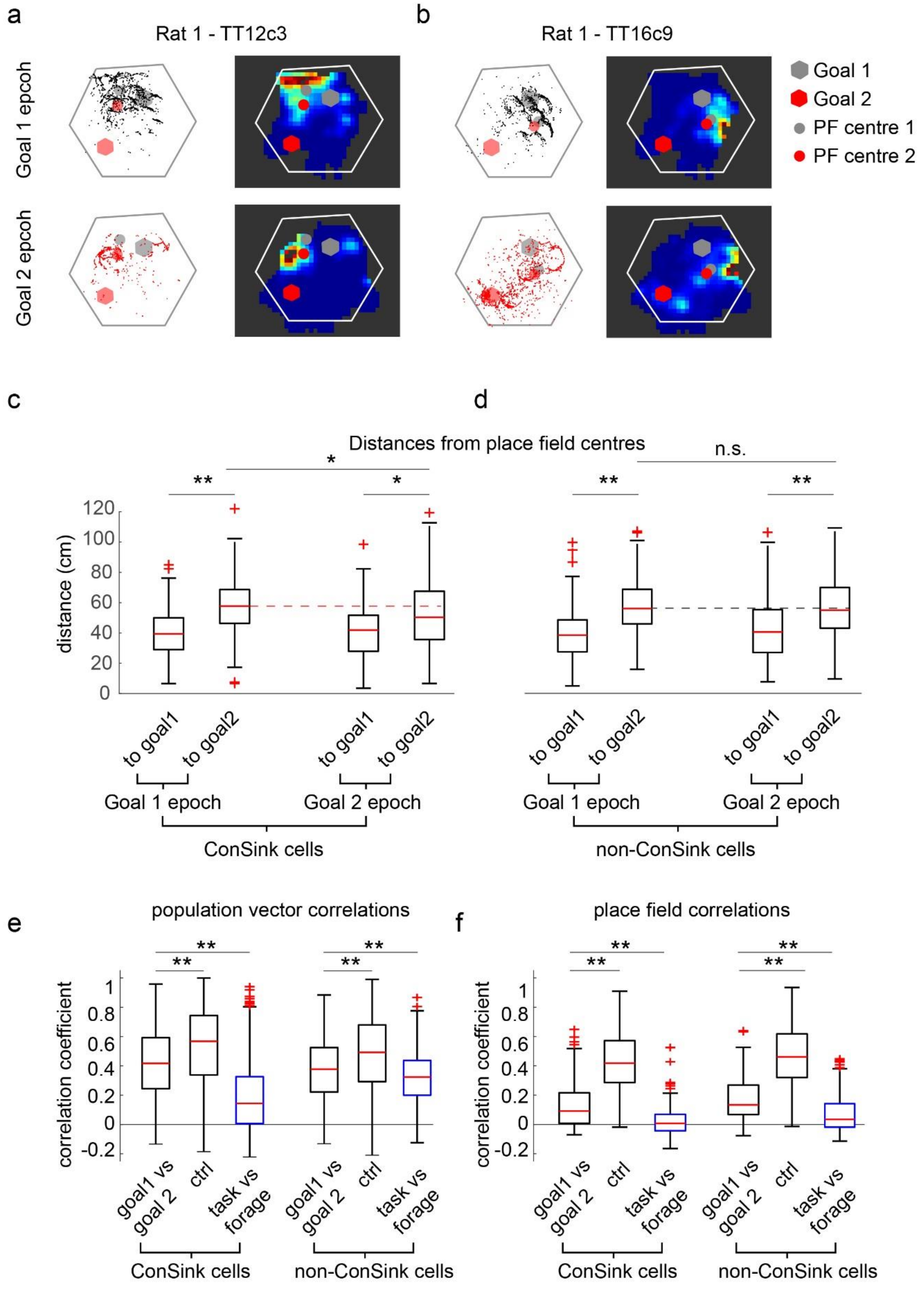
a
Rat 1 - TT12c3
b
Rat 1 - TT16c9
Goal 1 epcoh
Goal 2 epcoh
Goal 1
Goal 2
PF centre 1
PF centre 2
c
d
Distances from place field centres
*
**
*
n.s.
**
**
distance (cm)
0
20
40
60
80
100
120
to goal1
to goal2
Goal 1 epoch
Goal 2 epoch
ConSink cells
non-ConSink cells
e
population vector correlations
f
place field correlations
correlation coefficient
-0.2
0
0.2
0.4
0.6
0.8
1
goal1 vs goal 2
ctrl
task vs forage
ConSink cells
non-ConSink cells

**Extended Data Figure 9 │ Place fields do not show clustering around goal 2 after the goal switch. a,** Spike plots and rate maps from a representative relative direction cell during Goal 1 and Goal 2 epochs. Note that the rate map centre of mass (place field centre) shifts slightly towards Goal 2 after the goal switch, but remains closer to Goal 1. **b**, A second representative example cell. Again, centre of mass shifts slightly towards Goal 2 but remains closer to Goal 1. **c**, **d**, Summary of place field centre – goal distance data. Note that results are similar for both relative direction (**c**) and non-relative direction (**d**) cells; place field centres are initially closer to Goal 1 than to Goal 2, and remain so after the goal switch, though in relative direction cells only, they do move significantly towards Goal 2. Wilcoxon rank sum test, Goal 2 epoch – goal1 vs goal2, $p = 0.0017$; relative direction cells Goal 1 epoch – goal 2 vs Goal 2 epoch – goal 2, $p = 0.024$; non-relative direction cells Goal 1 epoch – goal 2 vs Goal 2 epoch – goal 2, $p = 0.59$; all other comparisons $p < 0.001$. **e**, **f**, Remapping in both relative direction (**e**) and non-relative direction (**f**) cells after the goal switch is less than is observed after the switch from honeycomb task to open field foraging. Remapping defined as significantly lower correlation than within control data; control data consists of session 1 (i.e. static goal) data split into first and second halves. Wilcoxon rank sum test. * indicates $p < 0.05$, ** indicates $p < 0.001$.

a

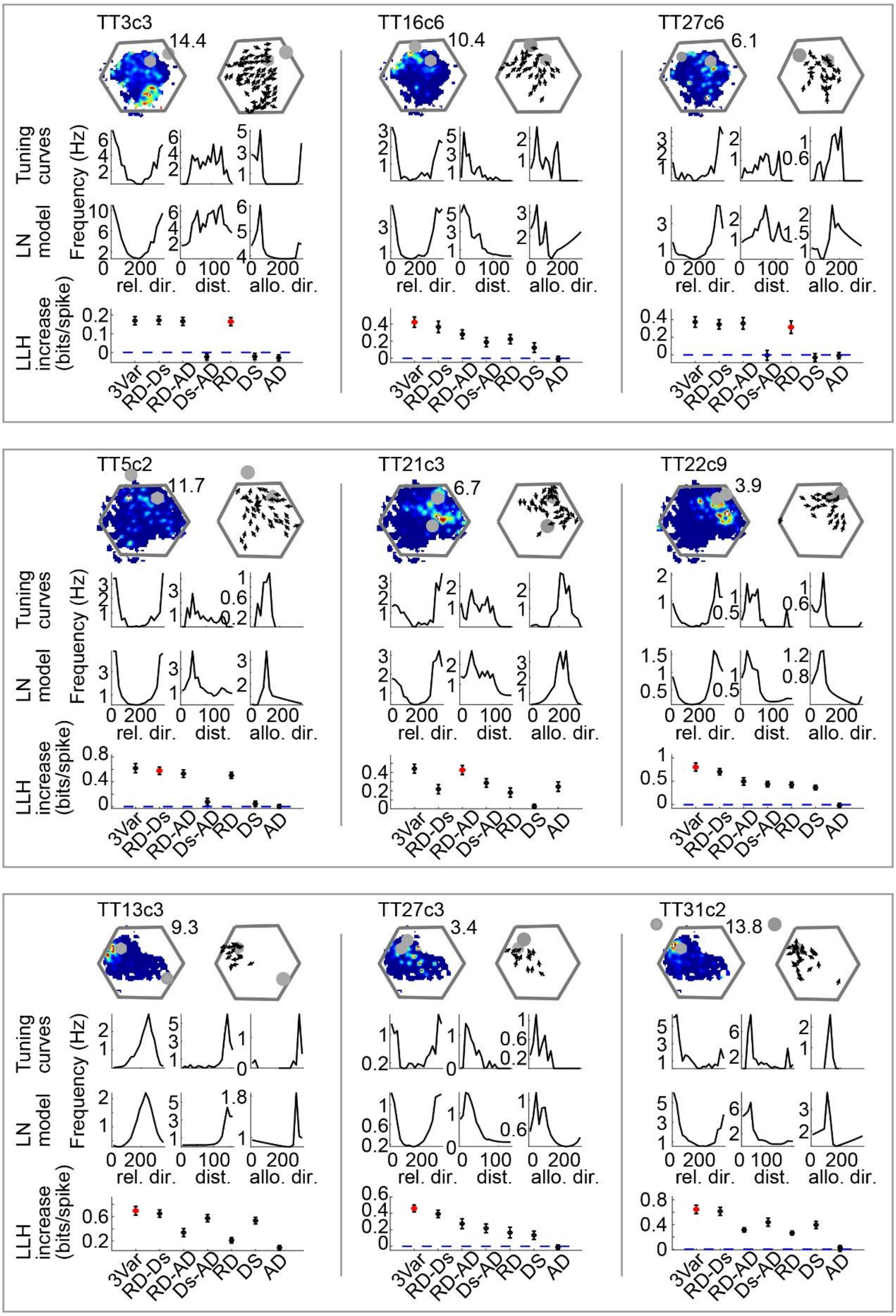
Rat 1
Rat 2
Rat 3
TT3c3
14.4
TT16c6
10.4
TT27c6
6.1
TT5c2
11.7
TT21c3
6.7
TT22c9
3.9
TT13c3
9.3
TT27c3
3.4
TT31c2
13.8
Tuning curves
LN model
Frequency (Hz)
rel. dir.
dist.
allo. dir.
LLH increase (bits/spike)
3Var
RD-Ds
RD-AD
Ds-AD
RD
DS
AD

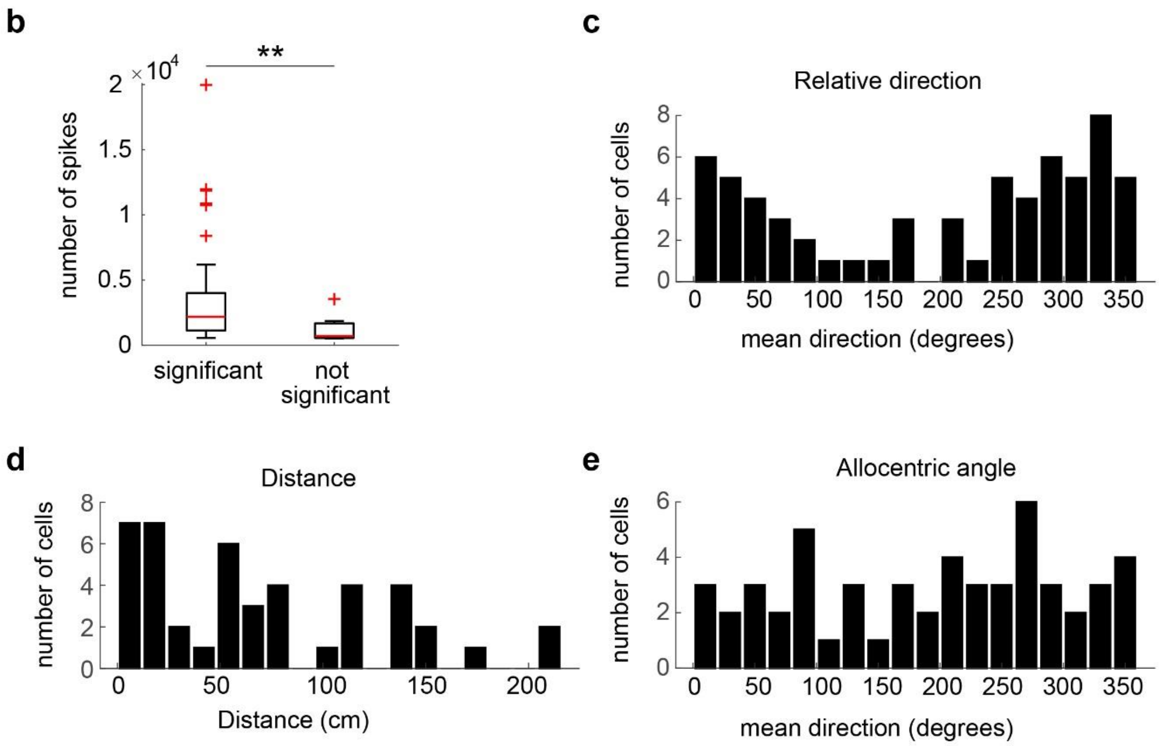


**Extended Data Figure 10 │ Representative examples of LN model results. a**, The LN model analysis found that ConSink cells significantly encoded various combinations of relative direction, allocentric direction, and distance to the sink. Plots for individual cells show each cell's rate map (top left) with the goal (grey hexagon) and ConSink (grey closed circle) and the peak firing rate (to the top right of the rate map; the vector field (top right) showing spike-associated allocentric head direction by spatial position; tuning curves for relative direction, distance, and allocentric direction to the sink (second row, from left to right); the LN model response profiles (third row); the log likelihood increase in information about the firing rate for each nested LN model (3VAr indicates the model incorporating all 3 variables; RD, relative direction; Ds, distance; AD, allocentric direction). The significant model (i.e. the highest order model that produces a significant increase in information over the next lower-ordered model) is indicated in red. **b**, Cells not found significant by the LN analysis fired fewer spikes than significant cells. Wilcoxon rank sum test, $p < 0.001$.

**c**, Relative direction, **d**, Distance, and **e**, Allocentric angle are well represented across the spectrum of possible values. Plots show the population of peak values taken from the subset of ConSink cells found significant for the respective spatial variable.

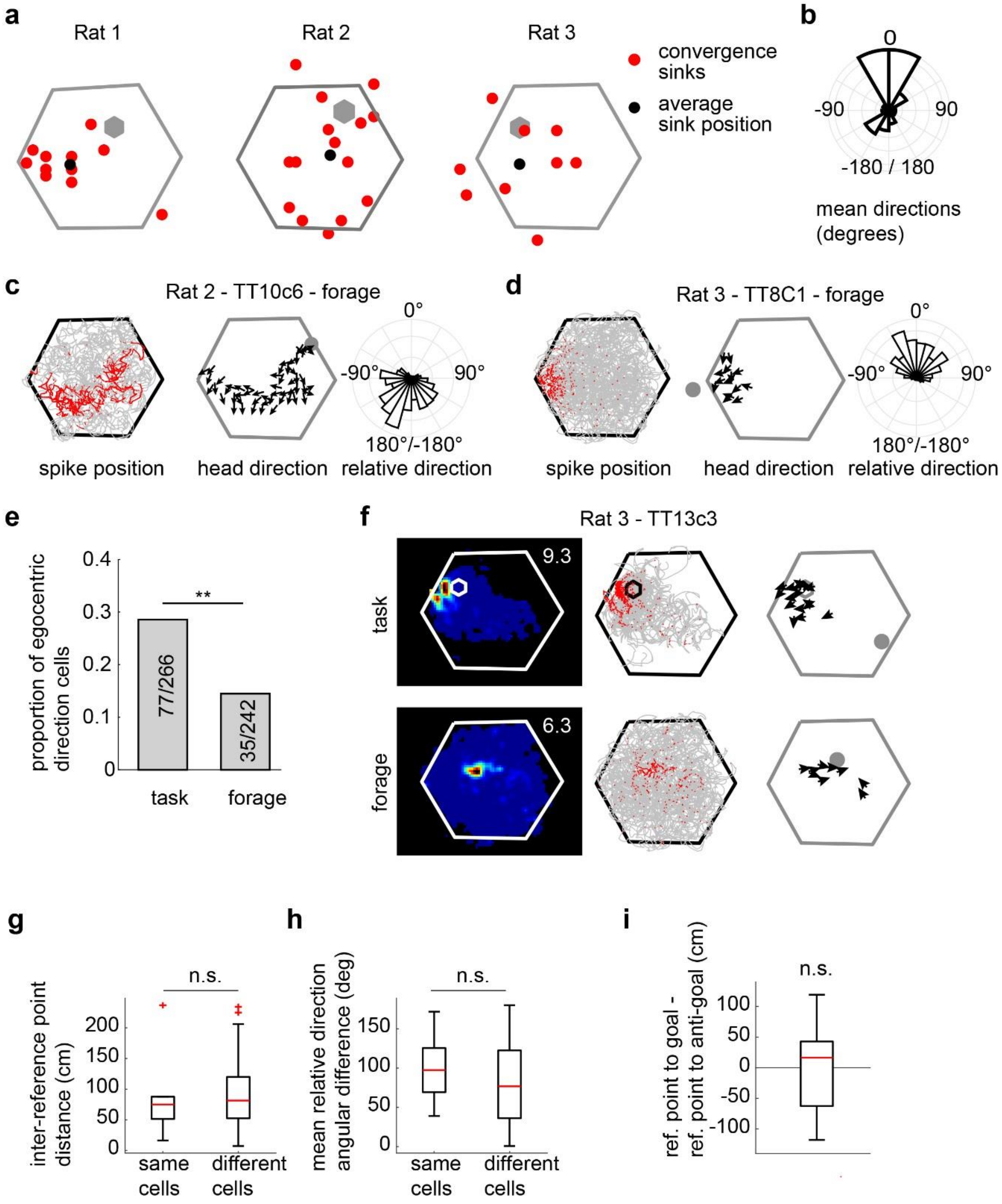

**Extended Data Figure 11 │ ConSinks during open field foraging are fewer and not clustered. a,** Distribution of ConSinks and average ConSink for each animal. **b**, Average heading direction relative to the ConSink across all animals. **c**, **d**, Representative examples of ConSink cells during foraging. Left, paths (grey) and spikes (red). Middle, vector field depicts mean head direction at binned spatial positions; ConSink is shown as grey filled circle. Right, Polar plot showing the distribution of spike-associated head directions relative to the ConSink. **e**, Twice as many place cells were significant for ConSink tuning during navigation compared to foraging ($p < 0.001$, Chi-square test). **f**, Example of a Ca1 place cell with significant ConSinks during both navigation and foraging; note differences in field location, vector fields, and ConSink location. Peak firing rate is indicated at top right of rate map (left-most panel). **g**, The convergence sinks for the task and forage tasks were no closer within cells that were significantly modulated during both tasks than they were across cells significantly modulated during either tasks, indicating a reorganization between the two epochs (n = 6 (same cells), 456 (different cells), Wilcoxon signed rank test, $p = 0.686$). **h,** The same lack of a difference held for the preferred relative directions ($p = 0.358$). **i,** ConSinks during foraging were no closer to the honeycomb task goals than to symmetrical locations on the opposite side of the maze (Wilcoxon signed rank test, $p = 0.623$; see Extended Data Fig. 5 for anti-goal positions).

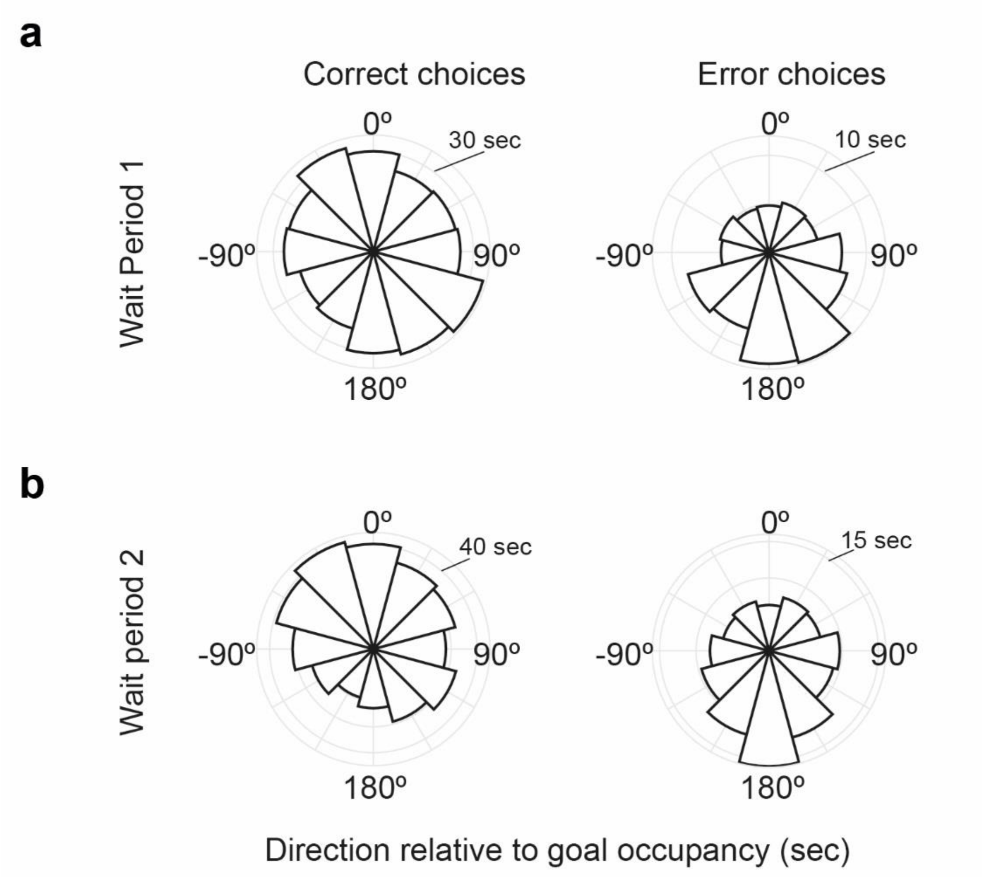


**Extended Data Figure 12 │ Behavioural orientations during correct and error choices. a**, The time spent oriented towards the goal in each possible orientation, averaged across the 3 animals, during Wait Period 1, which occurred after the previous platforms had lowered but before the next choice platforms were raised. Note the relatively uniform distribution during correct choices, while during errors, the animals orient away from goal (Rayleigh test for non-uniformity, Correct choices, $p = 0.78$, Error choices, $p = 0.015$). **b**, As in (**a**), but for Wait Period 2, which we define as the 4 seconds preceding the animals movement onto the Choice platform (Rayleigh test for non-uniformity, Correct choices, $p = 0.0014$, Error choices, $p = 0.022$).
